# Diverse 3D cellular patterns underlie the development of *Cardamine hirsuta* and *Arabidopsis thaliana* ovules

**DOI:** 10.1101/2023.12.06.570408

**Authors:** Tejasvinee Atul Mody, Alexander Rolle, Nico Stucki, Fabian Roll, Ulrich Bauer, Kay Schneitz

**Affiliations:** Plant Developmental Biology, TUM School of Life Sciences, Technical University of Munich, Emil-Ramann-Str. 4, 85354 Freising, Germany; Applied and Computational Topology, TUM School of Computation, Information and Technology, Technical University of Munich, Boltzmannstr. 3, 85747 Garching, Germany; Munich Data Science Institute, Technical University of Munich, Walther-von-Dyck Str. 10, 85747 Garching, Germany

**Keywords:** 3D digital organ, ovule, Cardamine, Arabidopsis, morphogenesis, evo-devo, topology

## Abstract

A fundamental question in biology is how organ morphogenesis comes about. The ovules of *Arabidopsis thaliana* have been established as a successful model to study numerous aspects of tissue morphogenesis; however, little is known regarding the relative contributions and dynamics of differential tissue and cellular growth and architecture in establishing ovule morphogenesis in different species. To address this issue, we generated a 3D digital atlas of *Cardamine hirsuta* ovule development with full cellular resolution. We combined quantitative comparative morphometrics and topological analysis to explore similarities and differences in the 3D cellular architectures underlying ovule development of the two species. We discovered that they show diversity in the way the three radial cell layers of the primordium contribute to its growth, in the formation of a new cell layer in the inner integument and, in certain cases, in the topological properties of the 3D cell architectures of homologous tissues despite their similar shape. Our work demonstrates the power of comparative 3D cellular morphometry and the importance of internal tissues and their cellular architecture in organ morphogenesis.

**Summary Statement:** Quantitative morphometric comparison of 3D digital ovules at full cellular resolution reveals diversity in internal 3D cellular architectures between similarly shaped ovules of *Cardamine hirsuta* and *Arabidopsis thaliana*.

## Introduction

Organisms exhibit a dazzling variety of species-specific sizes and shapes. Despite impressive progress, how an organ achieves its size and shape remains a major unresolved biological question. The last decades witnessed astounding advances in the understanding of the molecular mechanisms underlying pattern formation and morphogenesis. Much less well known, however, are the complex cellular behaviors that often lead to emergent tissue properties and ultimately to the functional architectures that make up a tissue or organ, although impressive progress has been made in dissecting the cellular basis of morphogenesis in, for example, epithelial tissues (Gómez-Gálvez et al., 2021; Lemke and Nelson, 2021; Sozen et al., 2022).

Plants are uniquely suited to study the cellular basis of tissue morphogenesis. Plant species are extremely diverse in size and shape, but are composed of relatively few cell types and tissues made up of immobile cells (Lyndon, 1990). To understand the differences in the 3D cellular architectures that underlie differences in morphogenesis, it is paramount to decipher the 3D cellular architecture of tissues at different developmental stages to enable quantitative 3D single cell analysis also in internal tissues. Advances in 3D confocal imaging of fixed, cleared, and stained organs, along with the development of open-source software for image processing, including 3D segmentation and mesh generation, have enabled 3D analysis at the cellular, tissue, and organ levels (Barbier de Reuille et al., 2015; Kurihara et al., 2015; Musielak et al., 2015; Strauss et al., 2019; Strauss et al., 2022; Tofanelli et al., 2019; Truernit et al., 2008). These advances allowed digital single cell analyses of organs with regular internal cellular architecture and simple shape, including early embryo, hypocotyl, or root (Bassel et al., 2014; Duran-Nebreda et al., 2023; Fridman et al., 2021; Graeff et al., 2021; Hernandez-Lagana et al., 2021; Lora et al., 2017; Montenegro-Johnson et al., 2015; Ouedraogo et al., 2023; Pasternak et al., 2017; Schmidt et al., 2014; Vijayan et al., 2021; Yoshida et al., 2014).

Most of these 3D digital single cell analyses have focused on organs of *Arabidopsis thaliana* (*A. thaliana)*. Comparative studies can explore the extent of cellular diversity underlying morphogenesis across species and possibly result in the emergence of a finite set of formal rules that enable cell collectives to organize into functional architectures. An evolutionary developmental (evo-devo) approach to understanding organogenesis in plants has been tremendously successful in the study of the genetic and molecular basis of floral organ identity, where comparative approaches have revealed much about the similarities and differences in this process across higher plants (Causier et al., 2010; Chanderbali et al., 2016; Coen and Meyerowitz, 1991; Kramer, 2019; Rümpler and Theißen, 2019). A similarly deep understanding of the 3D cellular basis of morphogenesis in different plants, especially with respect to internal tissues, is presently not available and is urgently needed. Thus, there is an evident requirement to extend quantitative 3D single cell studies of organogenesis to other species.

*Cardamine hirsuta* (*C. hirsuta*) represents an exciting model system for comparative analysis of plant morphogenesis (Hay and Tsiantis, 2016; Hay et al., 2014). The species displays abundant morphological diversity compared to its relatively recently diverged and reproductively isolated relative *A. thaliana*, providing a solid basis for investigating the evolutionary and ecological basis of morphological diversity. For example, it has been extensively used to investigate the genetic and cellular basis of the different leaf morphologies of the two species (Hay and Tsiantis, 2006; Kierzkowski et al., 2019; Vlad et al., 2014).

Ovules are elaborate-shaped organs that play a central role in female sexual reproduction in higher plants. At maturity, they carry the embryo sac with the actual egg cell enclosed by one or two integuments that protect the embryo sac and later develop into the seed coat. Several characteristics underlie ovule diversity among plant species, including ovule size, number of integuments, and degree of curvature (Endress, 2011). Ovule morphology has been extensively studied across a broad variety of plant species (Endress, 2011; Gasser and Skinner, 2019). However, a comprehensive quantitative understanding of the 3D cellular architecture underlying the differences in ovule morphology between species is lacking.

Ovules of *A. thaliana* along with their exciting biology, are an established successful model system to study different aspects of tissue morphogenesis (Chaudhary et al., 2018; Cucinotta et al., 2014; Gasser and Skinner, 2019; Nakajima, 2018). Recently, a reference 3D digital atlas of *A. thaliana* ovule development with full cellular resolution was established (Vijayan et al., 2021). The atlas enabled quantitative analysis of 3D cell and tissue growth patterns and dynamics even for an organ with an intricate internal cellular architecture and complex overall shape. It provides an excellent reference for a detailed 3D comparative analysis of ovule development. In terms of ovule morphology *C. hirsuta* and *A. thaliana* show differences in the shape of mature pre-fertilization ovules, as well as in ovule size, seed weight, size, and shape (Hay et al., 2014; Neumann and Hay, 2020). The general differences in ovule and seed architecture between *C. hirsuta* and *A. thaliana* suggested that a detailed comparative cellular analysis of ovule development would provide new insights into the 3D cellular architectures and the resulting morphogenetic differences of the ovules of the two species.

Here, we present a deep imaging-based reference 3D digital atlas of *C. hirsuta* ovule development with full cellular resolution. We further combined a comparative morphometric evo-devo approach with mathematical analysis of the topology of the 3D cellular architecture. Using this interdisciplinary strategy we identified and quantified similarities and differences in tissue growth and morphology and elucidated the role of the internal cellular architecture of tissues in shaping morphological differences between ovules of the two plant species.

## Results

### Generation of a reference 3D digital atlas of *C. hirsut*a ovule development with cellular resolution

To generate a comprehensive 3D digital atlas of *C. hirsuta* ovule development we followed the procedure outlined in (Tofanelli et al., 2019; Vijayan et al., 2021) and in the Materials and Methods section. In short, we performed 3D confocal laser scanning microscopy of fixed, cleared (Kurihara et al., 2015), and stained ovules to obtain z-stacks of optical sections of ovules of all stages (staging according to (Schneitz et al., 1995; Vijayan et al., 2021)). Ovules were stained with SR2200 to mark the cell outlines and TO-PRO-3 iodide to label the nuclei (Bink et al., 2001; Van Hooijdonk et al., 1994). This was followed by cell boundary prediction and 3D cell segmentation using the PlantSeg pipeline (Wolny et al., 2020). The segmented stacks were then loaded into MorphoGraphX (MGX) software for mesh generation, cell type labeling, and quantitative analysis. The entire procedure, from imaging to the final segmented and labeled 3D digital ovule, takes approximately 2 hours per z-stack. It is about twice the time required for *A. thaliana* ovules because *C. hirsuta* ovules are considerably larger in size. Overall, these efforts resulted in the generation of a high quality reference set of 144 hand-curated 3D digital ovules of the wild-type Oxford accession of *C. hirsuta* (≥10 samples per stage, 13 stages from stage 1-I to 3-VI) (Fig. 1). The entire *C. hirsuta* ovule dataset has been deposited with the BioStudies data repository (Sarkans et al., 2018) (https://www.ebi.ac.uk/biostudies) (accession number S-BIAD957).

**Fig 1.**
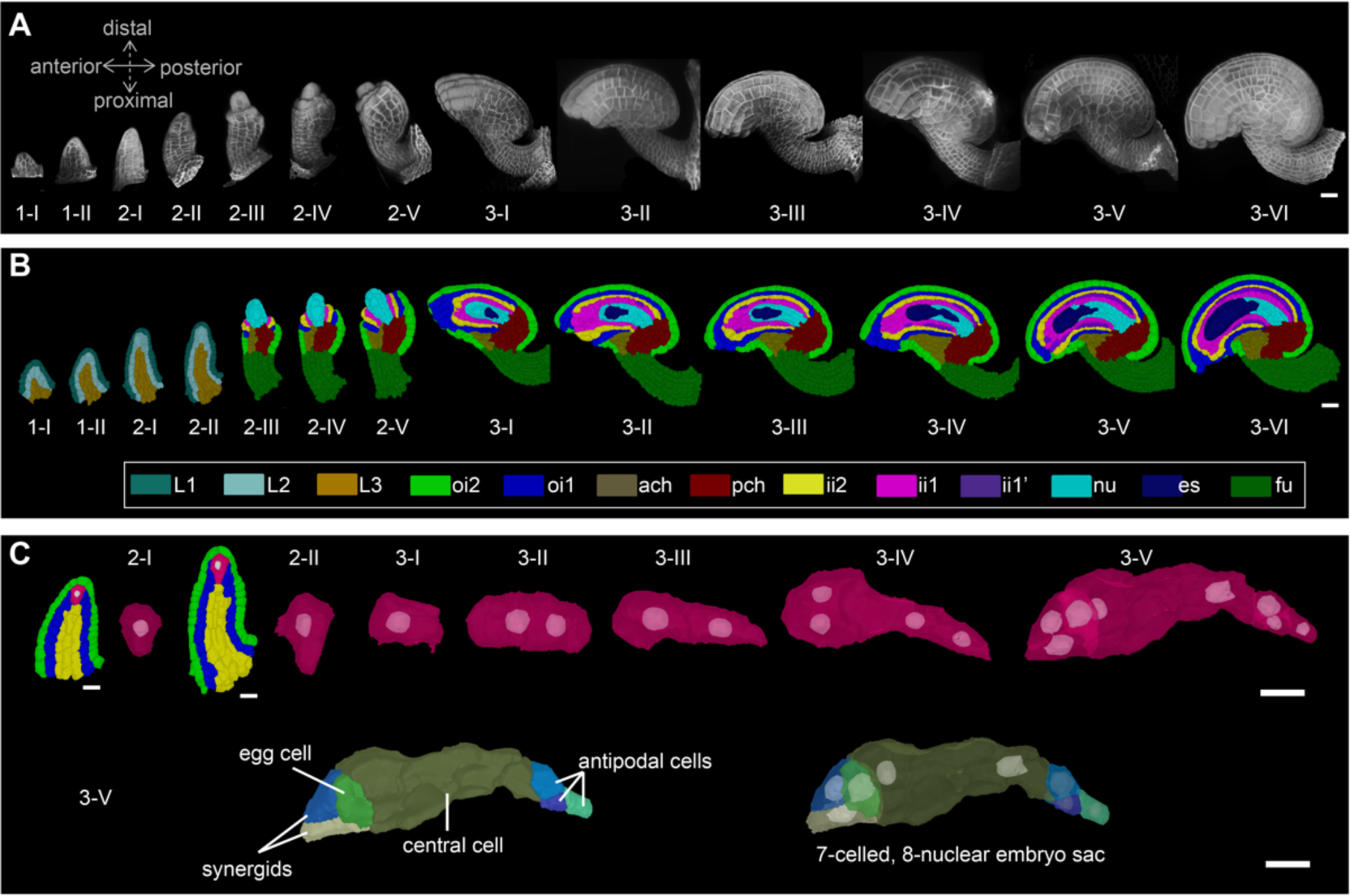
A 3D digital atlas of *C. hirsuta* ovule development. (A) *C. hirsuta* ovule development from initiation at stage 1-I to maturity at stage 3-VI. Images are 3D renderings of CLSM z-stacks of ovules with SR2200-stained cell walls. The anterior-posterior and proximal-distal axes are indicated. (B) Approximate mid-sagittal 2D sections of cell-type labeled 3D ovule meshes from stages 1-I to 3-VI showing the cell type organization in wild-type ovules. Stages 1-I to 2-II labeling includes radially organized L1, L2, L3 layers. From stage 2-III up to 3-VI, individual cell-type labels are tissue-specific and are represented with different colors. (C) Megaspore mother cell development in stage 2 primordia and embryo sac development from mono-nuclear embryo sac (stage 3-I) up to seven-celled, eight-nucleate embryo sac (stage 3-VI). Abbreviations: ach, anterior chalaza; es, embryo sac; fu, funiculus; ii1, inner layer of inner integument; ii1’, parenchymatic layer of inner integument; ii2, outer layer of inner integument; nu, nucellus; oi1, inner layer of outer integument; oi2, outer layer of outer integument; pch, posterior chalaza. Scale bars: A and B, 20 µm; C, 10 µm.

The regions and types of segmentation errors in the cell-segmented stacks of *C. hirsuta* ovules are similar to those observed in the case of *A. thaliana* ovule segmentation ((Tofanelli et al., 2019; Vijayan et al., 2021). The main sources of error include two groups of cells: the megaspore mother cell (MMC) and its immediate lateral neighbors at stages 2-II to 2-V form the first group and the embryo sac cells at stages 3-V and 3-VI form the second group. The main reasons for these errors could be insufficient cell wall staining with SR2200 or partially formed cell walls (Mansfield et al., 1991). Oversegmentation errors, if present, were corrected manually. Stage 1-I to 2-I and 3-I to 3-IV ovules selected for quantitative analysis had no apparent segmentation errors. Stage 2-II to 2-V ovules selected contained no more than five undersegmented (uncorrected) cells in the region occupied by the MMC and its lateral neighbors. Stage 3-V to 3-VI mature ovules selected had no obvious segmentation errors in the sporophytic tissue.

### General morphology of *C. hirsuta* ovule development

The ovules of *C. hirsuta* exhibit an overall shape and tissue organization similar but not identical to that of the ovules of *A. thaliana* (Fig. 1A-C) (Robinson-Beers et al., 1992; Schneitz et al., 1995; Vijayan et al., 2021). To stage *C. hirsuta* ovule development we adopted the staging system of *A. thaliana* (Schneitz et al., 1995; Vijayan et al., 2021). During stage 1, *C. hirsuta* ovule primordia emerge from the placenta as finger-like protrusions. During stage 2, three pattern elements can be distinguished along the proximal-distal axis (Fig. 1B): the proximal funiculus, which connects the ovule to the placenta; the central chalaza, from which an inner and outer integument emerge laterally; and the distal nucellus, containing the large MMC, which undergoes meiosis, with one meiotic product developing into the haploid multicellular female gametophyte or embryo sac (Fig. 1C). For most of its development, each integument is composed of two cell layers, an inner or adaxial layer and an outer or abaxial layer. In a sagittal section, each of these layers has a single cell thickness. Development of the inner integument begins in a ring-like fashion and ends as a tube, while early development of the outer integument is centered on the posterior side of the chalaza, but continues laterally to partially encircle the chalaza forming a hood-like structure. During stage 3, the two integuments grow asymmetrically over the nucellus, and eventually the ovule exhibits a pronounced curvature. However, the final curvature is not as advanced at the anterior side of the ovule as in *A. thaliana* and appears slightly more hunchbacked posteriorly. The inner integument eventually generates a third layer. At their distal end, the integuments form a cleft, the micropyle, through which the pollen tube reaches the embryo sac containing the egg cell to affect fertilization. The position of the micropyle therefore influences pollen tube entry and hence fertilization.

During stage 2, meiosis occurs in the large sub-epidermal MMC and results in four megaspores, three of which degenerate and only the chalazal megaspore continues development and gives rise to the haploid embryo sac (Fig. 1C). Its development starts during early stage 3. The mono-nuclear embryo sac undergoes three rounds of nuclear division resulting in a syncytium of eight nuclei followed by rapid cellularization. Prior to fertilization, the female gametophyte contains a distal or micropylar egg apparatus with two synergid cells and the egg cell, the two-nucleate central cell with a large vacuole, and three antipodal cells at the proximal or chalazal pole. Thus, the *C. hirsuta* embryo sac is of the typical eight-nucleate, seven-cell polygonum type (Fig. 1C).

### Overall dimensions of *C. hirsuta* and *A. thaliana* ovules

The availability of the *C. hirsuta* 3D digital ovule dataset together with the *A. thaliana* 3D digital atlas (Vijayan et al., 2021), the only other published dataset covering the entire ovule development, enabled us to investigate the similarities and diversity of 3D cellular architectures underlying ovule development between the two species. To this end we performed a quantitative comparative study. We first asked the basic questions of how much the ovule size of the two species differs and what is the cellular basis of the difference. We quantified the ovule size of *C. hirsuta* in terms of total volume and cell number and found an incremental increase in both volume and cell number for each successive stage of development (Fig. 2A-C, Table 1). At stage 3-VI, the average cell number per *C. hirsuta* ovule is about 2160 cells (2159.5 ± 355.2 (mean ± SD)) and the mean volume per ovule is about 6.6 × 10^5^ μm^3^ (6.6 × 10^5^ ± 1.18× 10^5^). By contrast, the *A. thaliana* ovule at stage 3-VI features about 1900 cells (1897.0 ± 179.9) and a volume of 5 × 10^5^ μm^3^ (4.9 × 10^5^ ± 7.2× 10^5^). Thus, the stage 3-VI *C. hirsuta* ovule is approximately 34% larger than the *A. thaliana* ovule and has 14% more cells. At all other developmental stages, *C. hirsuta* ovules (Table 1) are also significantly larger in volume and have somewhat higher cell numbers and cell volumes than corresponding *A. thaliana* ovules (Table 1, Table S1, and Table 1 in (Vijayan et al., 2021)). In conclusion, the increase in ovule size in *C. hirsuta* compared to *A. thaliana* is mainly due to a larger average cell size, with a smaller contribution from cell number.

**Fig 2.**
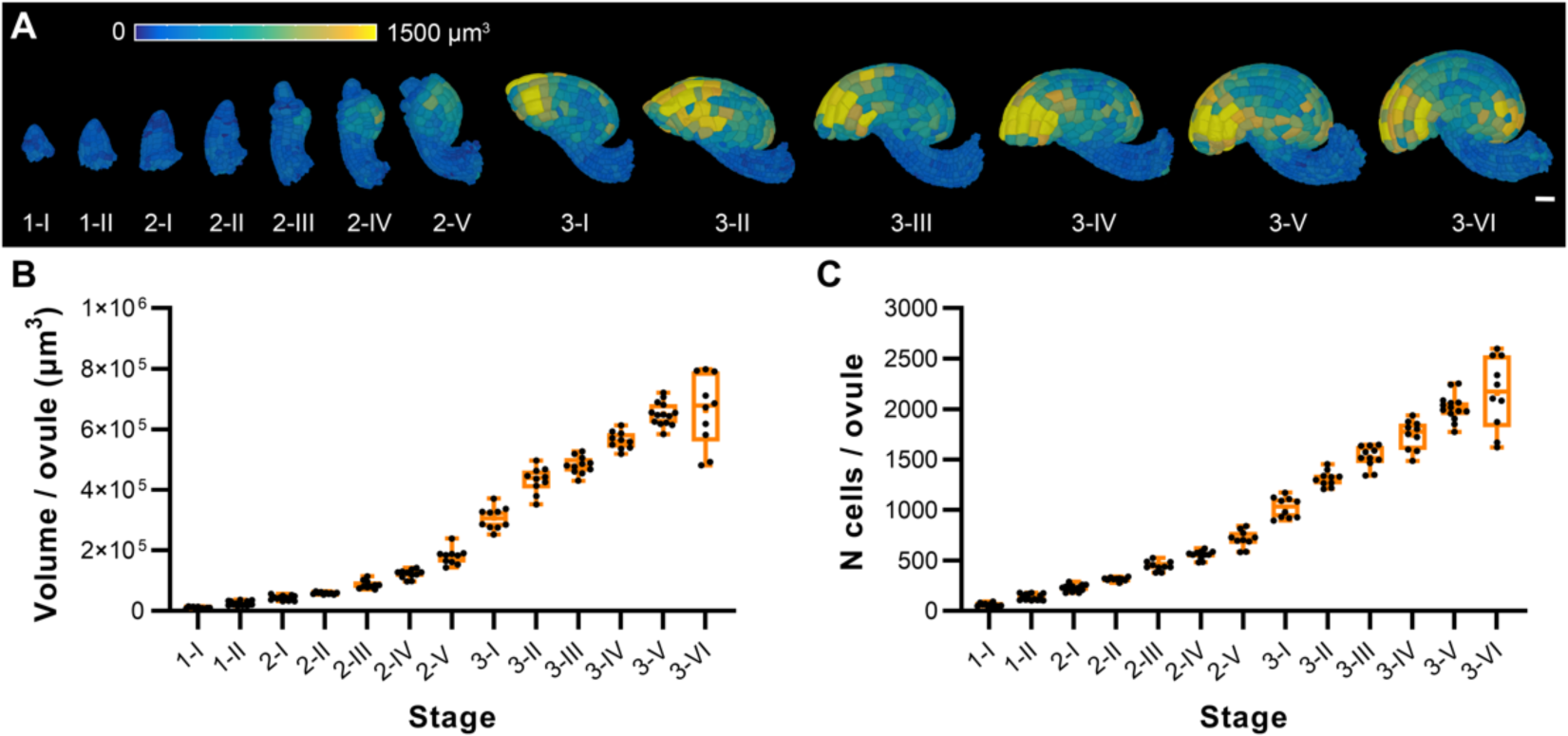
*C. hirsuta* overall growth patterns across ovule developmental stages. (A) 3D cell meshes of the developmental series of wild-type ovules, showing heat maps of the cell volume in the range from 0 to 1500 µm^3^. (B,C) Plots depicting the total volume and total number of cells of individual ovules from early to late stages of development, respectively. Data points indicate individual ovules. Mean ± SD is shown. Scale bar: 20 µm.

**Table 1.** Cell numbers, cell volumes, and total volumes of *C. hirsuta* ovules at different developmental stages.

| Stage | N<sup>a</sup> | N cells | Cell volume ($\mu\text{m}^3$) | Total volume ($\times 10^4 \mu\text{m}^3$) |
| --- | --- | --- | --- | --- |
| 1-I | 11 | 60.9 ± 16.4 | 170.1 ± 68.7 | 1.0 ± 0.3 |
| 1-II | 15 | 135.9 ± 31.1 | 181.5 ± 68 | 2.5 ± 0.7 |
| 2-I | 13 | 228.2 ± 33.2 | 185.0 ± 85.6 | 4.3 ± 0.8 |
| 2-II | 10 | 315.1 ± 17.4 | 184.9 ± 104.1 | 5.9 ± 0.4 |
| 2-III | 10 | 444.9 ± 45.8 | 194.6 ± 113.8 | 8.7 ± 1.3 |
| 2-IV | 11 | 555.0 ± 42.3 | 218.9 ± 137.4 | 12.2 ± 1.4 |
| 2-V | 10 | 714.1 ± 84.1 | 251.2 ± 184.1 | 18.0 ± 2.6 |
| 3-I | 10 | 1023.5 ± 101.7 | 297.0 ± 252.3 | 30.6 ± 3.6 |
| 3-II | 10 | 1311.5 ± 76.6 | 326.9 ± 297.4 | 43.2 ± 4.2 |
| 3-III | 11 | 1525.0 ± 107.8 | 312.8 ± 288.7 | 48.1 ± 2.8 |
| 3-IV | 10 | 1744.3 ± 144.6 | 318.0 ± 286.3 | 56.3 ± 2.9 |
| 3-V | 14 | 2014.3 ± 131.2 | 317.6 ± 294.4 | 65.1 ± 3.8 |
| 3-VI | 10 | 2159.5 ± 355.2 | 297.7 ± 253.8 | 66.3 ± 11.8 |

<sup>a</sup>Number of specimens scored.
Values represent mean ± SD.

### *C. hirsuta* and *A. thaliana* ovules exhibit tissue-specific growth patterns during development

Following tissue differentiation from stage 2-III onward, the different ovule tissues of *C. hirsuta,* with the exception of nucellus and embryo sac, tend to show larger total volumes, cell numbers and cell volumes than corresponding tissues of *A. thaliana* (Tables S2-4, Fig. S1, and Table 2 in (Vijayan et al., 2021)). Next, we asked whether there are tissue-specific differences in growth patterns during ovule development of *C. hirsuta* and *A. thaliana*, respectively, and if so, whether the two species display diversity in these growth patterns. To address this question, we performed a comparative analysis of stage-wise relative tissue growth, as well as tissue-specific cellular growth and proliferation of this process. We estimated the tissue-specific relative growth and cell proliferation during consecutive stages from stages 2-III up to stage 3-VI (Fig. 3A,B) as described in Materials and Methods. We observe that for both species, the outer integument grows the most between stages 2-III and 3-I, followed by a decline in relative growth in the following stages. From stage 2-III to 2-IV, the outer integument of *C. hirsuta* grows 1.4 times more than that of *A. thaliana.* Whereas, from 2-IV to 2-V, the outer integument of *A. thaliana* grows 1.6 times more than that of *C. hirsuta.* This relative growth is also evident when plotting the relative contribution of outer integument to the total ovule volume (Fig. S2). The relative outer integument proportion continues to increase until stage 3-I, and then stays relatively constant. From stage 2-III to 2-IV, the inner integument of *A. thaliana* grows 20% more than that of *C. hirsuta.* In the subsequent stages until 3-IV, the relative inner integument growth of both species is comparable. However, there are between-species differences in the extent of relative growth between consecutive stages. The relative tissue cell proliferation patterns do not fully explain the relative tissue growth patterns, suggesting that cell growth might play a role. In cases where the relative cell proliferation is lower than the relative tissue growth, we observe an increase in average cell volumes. For instance, from stage 3-I to 3-II the *C. hirsuta* inner integument grows by 84.3 %, whereas there is a 55.4 % increase in tissue cell proliferation and 14% increase in average cell volume. The chalaza of both species shows a consistently higher relative tissue growth than relative tissue cell proliferation along with a consistent increase in average cell volumes over development (Fig. 3A,B; Table S4). However, interestingly, the relative chalaza proportion is higher in *C. hirsuta* than in *A. thaliana* (Fig. S2). Therefore, the *C. hirsuta* chalaza is disproportionately larger than that of *A. thaliana*.

**Fig 3.**
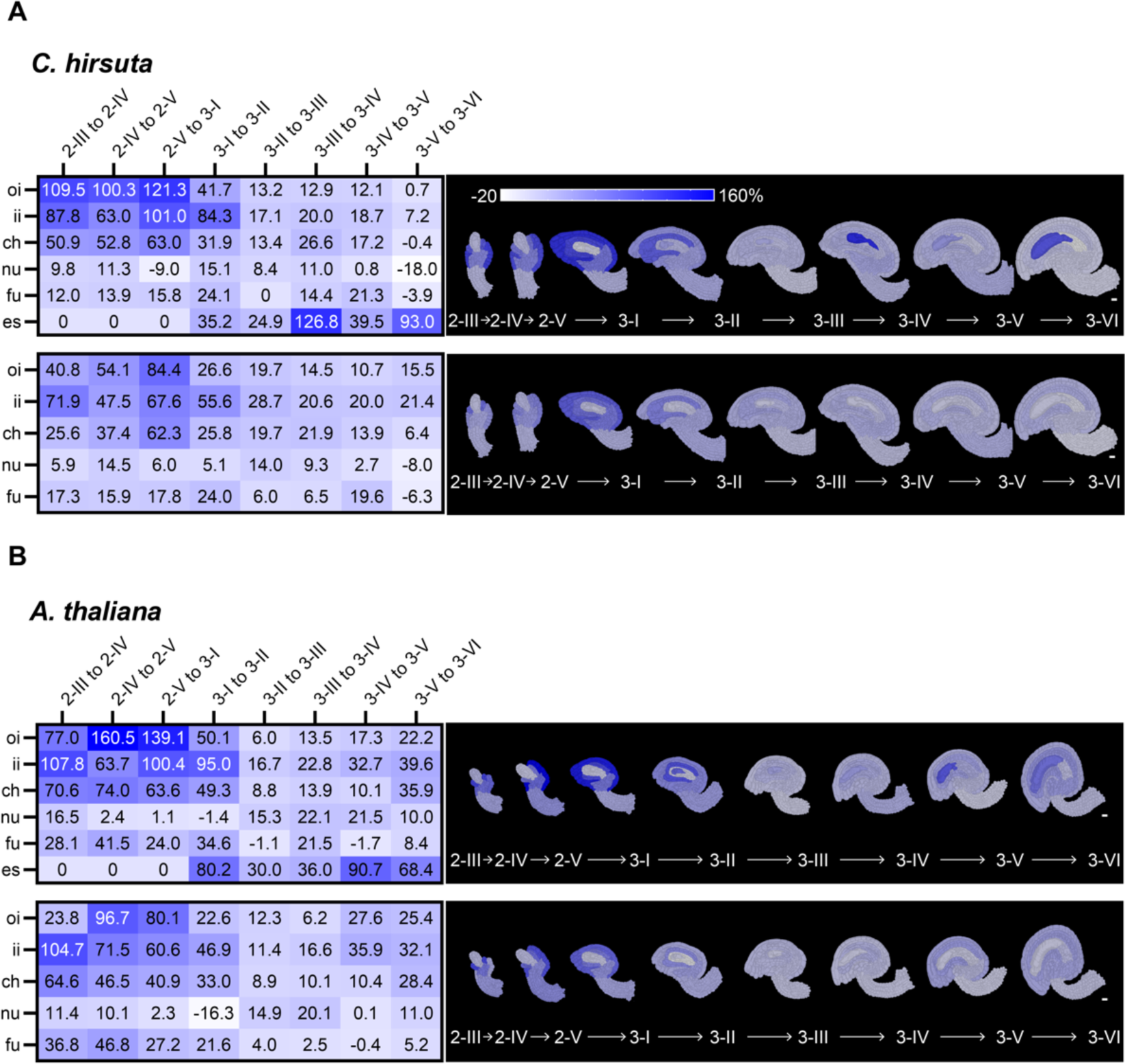
Comparative trends of tissue growth dynamics underlying *C. hirsuta* and *A. thaliana* ovule development. (A) *C. hirsuta* and (B) *A. thaliana* relative tissue growth (top) and relative tissue cell proliferation (bottom) between two consecutive stages. (A–B) Heatmaps (left) and mid-sagittal sections (right) depicting relative tissue growth (top) and cell proliferation (bottom) across the different ovule stages from 2-III to 3-VI. Heatmap values indicate % change in mean parameter from previous stage to the next relative to the previous stage. Scale bars: 10 µm.

**Table 2.**
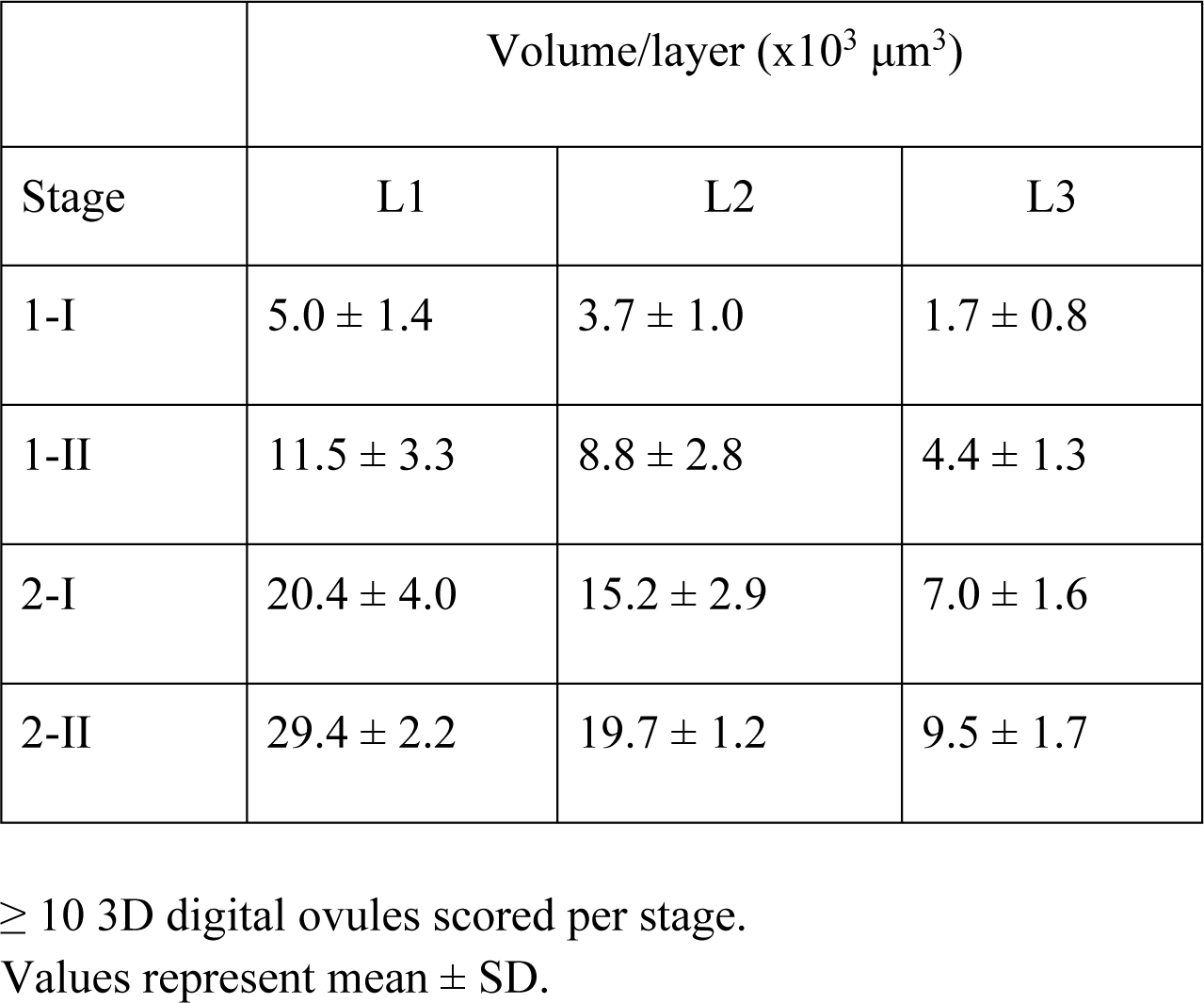
Layer volumes of *C. hirsuta* ovule primordia.

**Table 3.** Cell numbers in the different radial layers of *C. hirsuta* ovule primordia.

|  | Cell number/layer |  |  |
| --- | --- | --- | --- |
| Stage | L1 | L2 | L3 |
| 1-I | 34.8 ± 8.5 | 18 ± 5.0 | 8.1 ± 3.7 |
| 1-II | 70.5 ± 16.2 | 42.7 ± 10.1 | 23.1 ± 6.3 |
| 2-I | 118.4 ± 17.4 | 71.5 ± 11.1 | 38.3 ± 7.2 |
| 2-II | 162.4 ± 7.5 | 94.9 ± 6.9 | 55.4 ± 7.7 |

≥ 10 3D digital ovules scored per stage. Values represent mean ± SD.

**Table 4.** Cell volumes in the different radial layers of *C. hirsuta* ovule primordia.

|  | Layer-specific cell volume (μm<sup>3</sup>) |  |  |
| --- | --- | --- | --- |
| Stage | L1 | L2 | L3 |
| 1-I | 159 ± 61.7 | 229 ± 81.9 | 229 ± 70.2 |
| 1-II | 165 ± 56.3 | 208 ± 72.1 | 192 ± 69 |
| 2-I | 173 ± 62.9 | 208 ± 86.5 | 183 ± 68.8 |
| 2-II | 181 ± 64.7 | 199 ± 98 | 171 ± 64.9 |

≥ 10 3D digital ovules scored per stage. Values represent mean ± SD.

In summary, the analysis of tissue-specific growth patterns of the two species has revealed stage-specific differences in relative tissue growth and enabled us to dissect the contributions of relative tissue cell proliferation and cell volumes to this process.

### Topological analysis reveals differences in 3D cellular architecture between *C. hirsuta* and *A. thaliana*

Assessment of relative tissue growth between successive developmental stages in the two species revealed interesting differences, particularly when comparing the chalaza and integuments. We next asked whether there are differences between species in the 3D cellular architecture of tissues during development.

To address this question, we performed a topological analysis of the 3D cellular architecture of the ovule tissues using the nerve construction, a standard tool in topology (Edelsbrunner and Harer, 2010) (Materials and Methods). In various fields of mathematics, the nerve is used to encode how a complex geometric object is assembled from simple pieces. Applied to a 3D digital ovule, the nerve captures detailed information about the cellular architecture of the ovule, or of its tissues. To evaluate the between-species differences in the nerves we computed, we applied a statistical two-sample test to the “feature vectors” of nerves, since standard statistical tests cannot be applied directly to sets of nerves. Feature vectors are vector representations of the nerves that can be analyzed statistically. The neighborhood of a cell in a 3D digital ovule refers to the collection of all cells that intersect with that cell. The feature vector of an ovule (or of one of its tissues) contains structural information about the neighborhood of each cell in that ovule (or in that tissue).

We investigated the 3D cellular architecture of the integuments, chalaza, nucellus and funiculus of *A. thaliana* and *C. hirsuta* from stage 2-III to 3-VI. We asked whether the information about the 3D cellular architecture contained in the nerve can distinguish between the tissues of these two species. For each developmental stage from 2-III to 3-VI we obtained two sets of feature vectors: one set representing the *C. hirsuta* tissue, and one set representing *A. thaliana* tissue. The feature vectors after applying principal component analysis (PCA) to stage 3-VI ovules and tissues of *C. hirsuta* and *A. thaliana*, taking into account all cells are shown in Fig. 4B-G (right panels). We applied the multivariate two-sample test of Baringhaus and Franz to these feature vectors (Baringhaus and Franz, 2004), and obtained p values via bootstrapping (Fig. 4C-G and Table S5). We also compared the 3D cellular architecture of the entire ovule of *A. thaliana* and *C. hirsuta* (Fig. 4B and Table S5). The p values indicate that there is evidence for a difference in the cellular architecture of the outer integument, inner integument, and chalaza between the two species, starting around stage 3-I for the outer integument and chalaza, and around 3-III for the inner integument (Fig. 4C-G and Table S5). Interestingly, the p values indicate evidence for a between-species difference in the cellular architecture of the entire ovule, except at stage 3-IV. As a positive control, we tested whether the nerve construction could distinguish between the outer integument and the chalaza in these two species. For both species and all developmental stages, the resulting p-value was at most 0.0001. This further underscores the importance of our topological approach for distinguishing the 3D cellular architecture of tissues.

**Fig 4.**
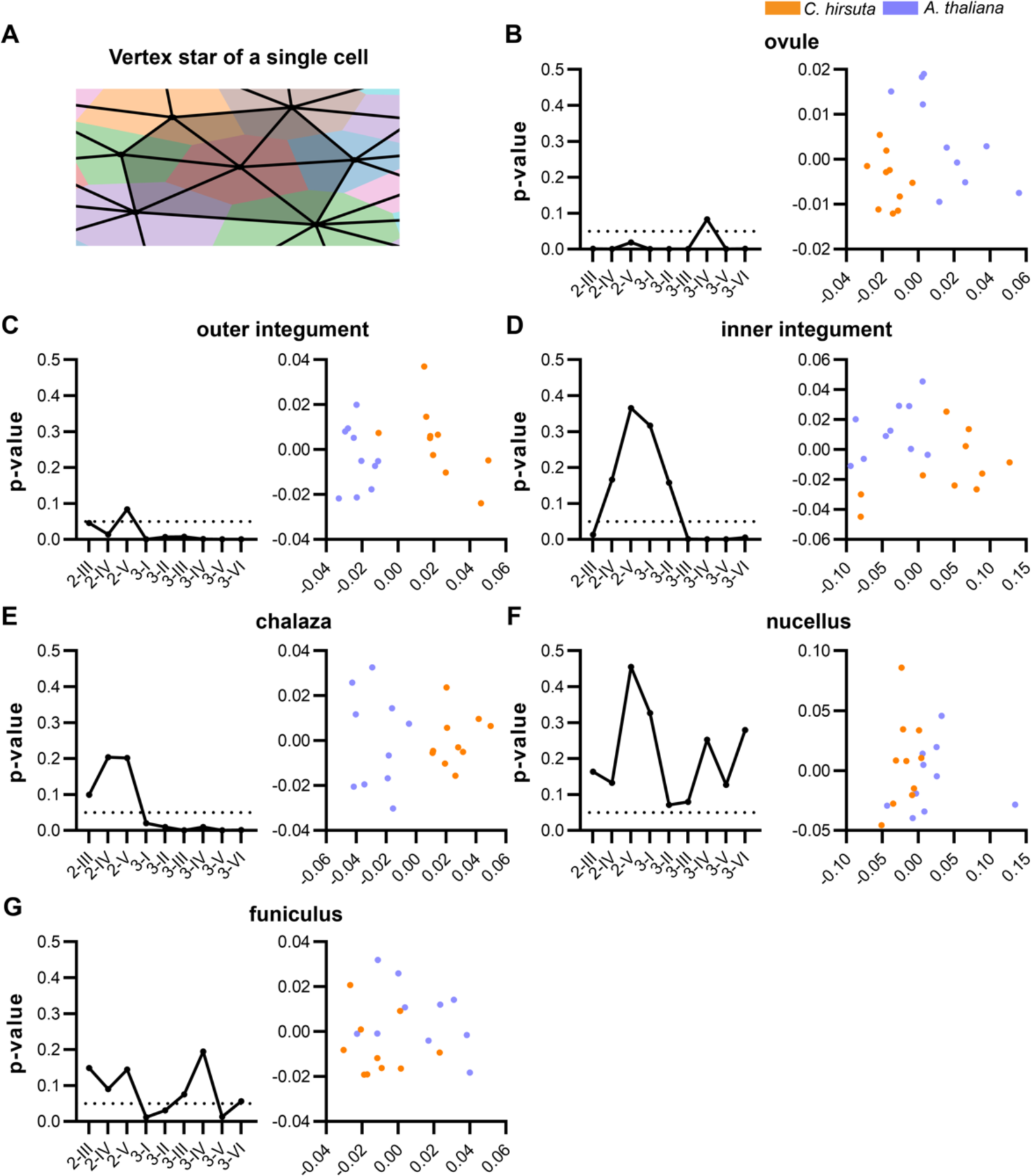
Topological analysis of 3D cellular architectures for *C. hirsuta* and *A. thaliana*. (A) The nerve complex (black lines) of a collection of cells (colored). The 0- and 1-simplices and the vertex star (1 vertex, 6 edges, 6 triangles; grey area) of the red cell are visualized. (B-G) Left panels: p-values obtained from comparison of *C. hirsuta vs A. thaliana* ovules across different stages, taking into account all cells of the ovule (B), only cells of outer integument (C), inner integument (D), chalaza (E), nucellus (F), and funiculus (G). The dashed line indicates p = 0.05. (B-G) Right panels: scatter plots of the feature vectors after applying PCA to stage 3-VI ovules of *C. hirsuta* (orange) and *A. thaliana* (blue), taking into account all cells of the analyzed ovule or tissue.

Together, these results suggest that the comparative differences in tissue growth are not merely related to differences in growth and proliferation at the cellular level, but may also be accompanied by differences in the cellular topology of tissues. Therefore, tissue growth occurs not only due to cell growth and proliferation, but also involves changes in 3D cellular architecture. These between-species differences in tissue growth and architecture result in the subtle differences in ovule morphology between the two species.

### The radial tissue layers of *C. hirsuta* and *A. thaliana* ovules contribute differently to ovule primordium outgrowth

We then focused our comparative analysis on selected developmental processes. We first examined primordium formation. The *A. thaliana* ovule follows the general principle of plant organ structure (Satina et al., 1940) and is characterized by three clonally distinct radial layers, the epidermal L1 layer, the sub-epidermal L2 layer, and the innermost L3 layer (Jenik and Irish, 2000). Three regular radial cell layers are readily recognizable in the ovule primordium of *C. hirsuta* suggesting a similar conventional tissue organization (Fig. 1B,C).

The onset of expression of a transcriptional reporter for the *WUSCHEL* (*WUS*) gene (pWUS) in *A. thaliana* primordia has been reported as a convenient marker to delineate non-pWUS-expressing stage 1-I primordia (< 50 cells) from pWUS-expressing stage 1-II primordia (> 50 cells) (Vijayan et al., 2021). The distinction between stage 1-II and stage 2-I in *A. thaliana* relies on the absence or presence of MMC formation with stage 2-I primordia being characterized by the presence of the MMC (Schneitz et al., 1995). Therefore, in *C. hirsuta*, we also distinguished stage 1 ovules from stage 2-I ovules based on the absence or presence of the MMC. The upper limit of total volume and cell number of *C. hirsuta* stage 1-II primordia was set based on the similarity in volume to stage 2-I primordia but absence of MMC. In *C. hirsuta*, in the absence of a pWUS reporter, we extrapolated the *A. thaliana* stage 1-II minimum and maximum values of total volume and cell number to estimate the lower limit of stage 1-II primordia to discriminate stage 1-II from stage 1-I. We used the formula [min_1-II, *C. hi.*_ = (min_1-II, *A. th.*_*/*max_1-II, *A. th.*_)*\** max_1-II, *C. hi*_] to set an approximate lower limit of total volume and cell numbers for stage 1-II *C. hirsuta* primordia.

Given these considerations, we defined stage 1-I of *C. hirsuta* ovules to include primordia containing up to 100 cells. The maximum volume of stage 1-I primordia is about 1.5 x 10^4^ µm^3^. The ovule primordia of stage 1-II have a maximum volume of up to 3.7 x 10^4^ µm^3^ and cell number range from 103-180. Stage 2-I primordia, as evidenced by the presence of a MMC, have a volume range of 3.2 x 10^4^ - 5.3 x 10^4^ µm^3^ and cell number range from 179-288. We found 5 out of 15 stage 1-II ovules and 3 out of 13 stage 2-I ovules with a cell number around 180 and a volume range of about 3.2 x 10^4^ - 3.7 x 10^4^ µm^3^. These results indicate that the transition from stage 1-II to 2-I in *C. hirsuta* occurs once the primordium has reached about 180 cells and a volume of about 3.5 x 10^4^ µm^3^. Whereas in *A. thaliana* the transition from stage 1-II to 2-I occurs at around 130 cells and a primordium volume of about 1.8 x 10^4^ µm^3^ (Vijayan et al., 2021).

The ovule primordium outgrowth in *C. hirsuta,* as in *A. thaliana*, is characterized by a steady and continuous increase in primordium volume and cell number until late stage 2-I (Fig. S3A). In addition, the *C. hirsuta* primordia show slanting (Fig. S3B), the first morphological manifestation of an anterior-posterior axis, as observed in *A. thaliana* (Vijayan et al., 2021). Despite the continuous growth, the transition from stage 1-II to 2-I seems to have a sharp 180-cell number cut-off. However, the primordia volume range of both these stages show an overlap between 3.2 x 10^4^ - 3.7 x 10^4^ µm^3^. This is also indicative of the primordium volume range at which MMC differentiation occurs. Early stage 1-II ovules that have a total volume smaller than that of the overlapping volume range, show a mean cell volume of 172.2 ± 63.5 µm^3^. Within this overlapping total volume range, the late stage 1-II ovules show a mean cell volume of 193.9 ± 71.8 µm^3^, whereas the early stage 2-I ovules have a mean cell volume of 175.1 ± 67.8 µm^3^.

Late stage 2-I ovules that have a total volume larger than that of the overlapping volume range, show a mean cell volume of 186.3 ± 77.9 µm^3^. Therefore, within both stages 1-II and 2-I the early to late transition involves an increase in cell volume. But in the progression from late stage 1-II to early 2-I there is a transient decline in cell size by about 10%. Therefore, despite increasing cell numbers during this transition, there is an overlap in total volumes of late stage 1-II and early stage 2-I ovules.

We then investigated whether the overall similarity in the outgrowth of the ovule primordia and the resulting cone-like shapes was reflected at the cellular and tissue level. To address this question, we performed a comparative analysis of stage-wise relative layer growth, as well as layer-specific cellular growth and proliferation of this process. We estimated the layer-specific relative growth during a given stage, up to stage 2-II, by applying the same formula to the various parameters as described above ((y(n + 1) - y(n))/y(n)). Unexpectedly, we observed surprising diversity in relative layer growth between *C. hirsuta* and *A. thaliana* (Fig. 5). Regarding the stage-wise growth of the individual layers, we found that in *C. hirsuta* most relative layer growth occurs during stage 1-I to 1-II, with steadily decreasing layer-specific relative growth during transition from stages 1-II to 2-I and 2-I to 2-II (Fig. 5A). Although we observed slight unequal relative growth of the individual layers the differences were relatively small. A markedly contrasting picture emerged when we examined the relative layer growth in *A. thaliana* ovule primordia (Fig. 5B). During the transition from stage 1-I to 1-II the L3 grew 2.6 to 3.8 times more than the L2 and L1, respectively. During stage 1-II to 2-I all layers grew by roughly similar percentages. From stage 2-I to 2-II, the L1 grew by 10%, while the L3 exhibited minimal growth. Similar scenarios were observed when comparing stage-specific relative cell numbers, although in *A. thaliana* the relative increase in cell numbers during stage 2-I to 2-II was comparable among the three layers and was accompanied by a significant decrease in mean cell volume in all three layers (Fig. 5A,B). In terms of the relative contribution of each layer to the total primordia volume and cell numbers, the L1 > L2 > L3 in both species. However, the L3 contribution is higher in *C. hirsuta,* whereas, the L1 contribution is higher in *A. thaliana* (Fig. 5C,D)

**Fig 5.**
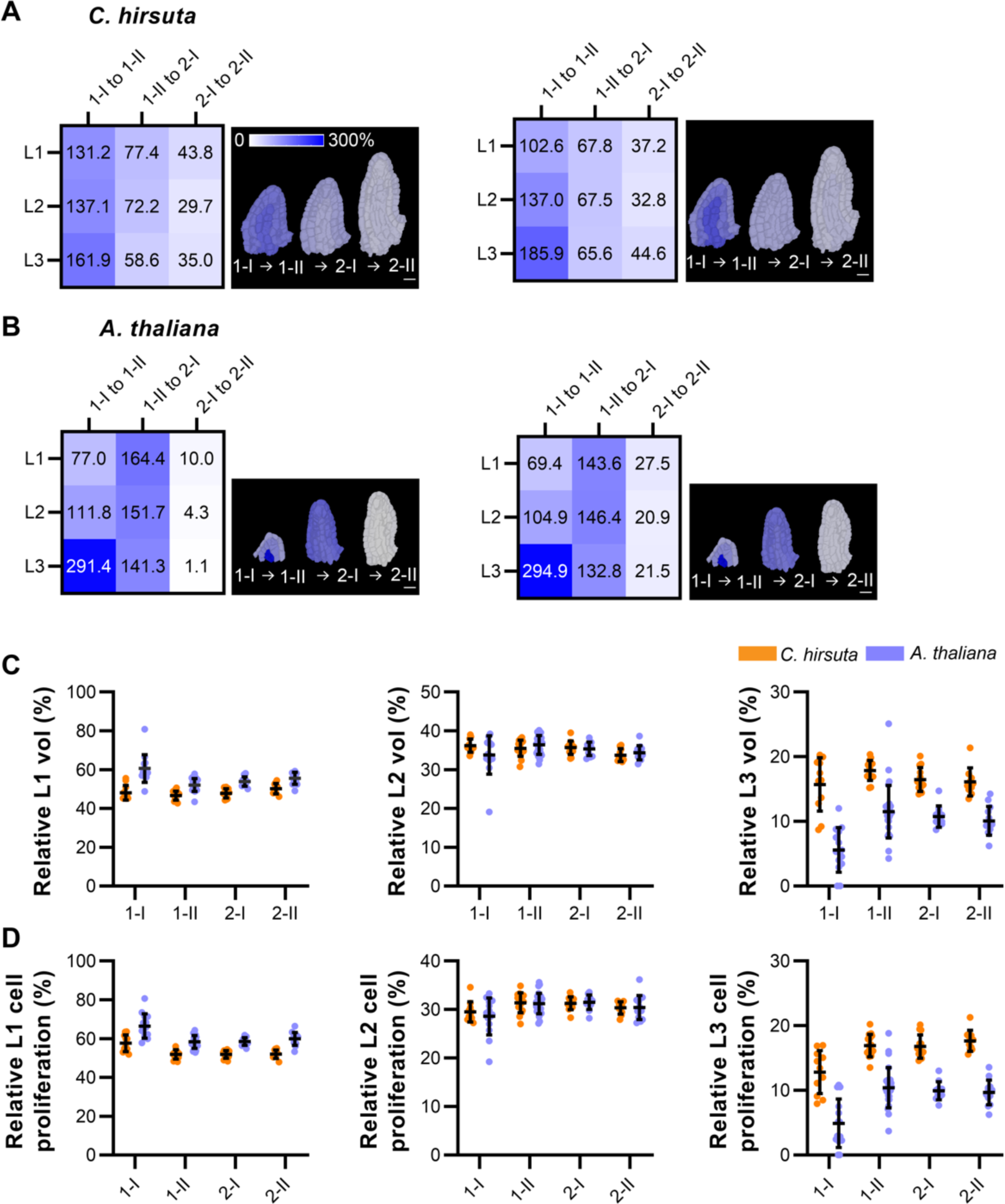
Differential radial layer-specific growth dynamics underlie *C. hirsuta* and *A. thaliana* early ovule development. (A) *C. hirsuta* and (B) *A. thaliana* relative tissue growth (left) and relative tissue cell proliferation (right) between two consecutive stages. (A,B) Heatmaps and mid-sagittal sections depicting relative tissue growth (left) and cell proliferation (right) across different consecutive ovule stages from 1-I to 2-II. Heatmap values indicate % change in mean parameter from previous stage to the next relative to the previous stage. (C,D) Plots depicting relative contribution of each radial tissue layer to the entire ovule primordium in terms of (C) total volume and (D) total cell proliferation. Data points indicate individual ovule primordia. Mean ± SD is shown. Scale bars: 10 µm.

We further compared early MMC development between species. The nucellus of *C. hirsuta*, like that of *A. thaliana* is tenuinucellate, that is, an archesporial cell in the first subepidermal cell layer directly develops into the MMC (Fig. 1C) (Dahlgren, 1927; van Tieghem, 1898). This is not true for all Cardamine species, e.g. the nucellus of *C. parviflora* is crassinucellate (Harvey and Smith, 2013), as an additional cell is located between the MMC and the L1, indicating that an archesporial cell has undergone a periclinal cell division resulting in a primary parietal and sporogenous cell, respectively, the latter of which developed into the MMC (Fig. S3C).

Interestingly, we observed a difference in MMC size regulation between *C. hirsuta* and *A. thaliana* (Fig. S3D). The *A. thaliana* MMC does not change its size between stages 2-I and 2-II, whereas the *C. hirsuta* MMC grows significantly between stages 2-I and 2-II, becoming even larger than that of *A. thaliana*. This subtle variation suggests a differential control of MMC size. However, the discrepancy in MMC size regulation does not lead to differences in the sizes of embryo sacs, as the dimensions of the mononuclear embryo sac (stage 3-I) and the final seven-cell embryo sac (stage 3-V) do not significantly differ between the two species (Table S2, Fig. S1, Figure 1-Appendix 1 in (Vijayan et al., 2021)).

In summary, the comparisons reveal similarities but also differences in the cellular basis of ovule primordium outgrowth between *C. hirsuta* and *A. thaliana*. First, they suggest that cell proliferation, rather than cell growth, predominantly underlies tissue growth patterns during early ovule development in both species. Second, they indicate that in each case the increase in primordium volume occurs in a steady and continuous fashion. Importantly, these results also show that despite similar overall primordium outgrowth and final primordium shape (and with the exception of size), the internal growth processes underlying ovule primordium outgrowth differ substantially between the two species.

### *C. hirsuta* and *A. thaliana* differ in the formation of the parenchymatic cell layer in the inner integument

Starting at stage 3-II, cambium-like activity of the inner (adaxial) layer of the inner integument (ii1, differentiating into the endothelium) in *A. thaliana* generates an additional cell layer (ii1’) located between the ii1 and ii2 layers of the inner integument (Debeaujon et al., 2003; Schneitz et al., 1995; Vijayan et al., 2021). Once formed, the ii1’ layer neither expresses the epidermis-specific *ARABIDOPSIS THALIANA MERISTEM L1* (*ATML1*) nor produces tannins like the ii1/endothelium layer, but remains parenchymal. In *A. thaliana* the ii1’ layer is generated by periclinal cell divisions of only a few scattered ii1 founder cells, followed by anticlinal cell divisions in the ii1’ daughter cells. Eventually, the ii1’ layer forms a ring-like structure covering roughly the proximal half of the inner integument. Formation of this new cell layer in *A. thaliana* exhibits features of layer invasion.

We could also observe the ii1’ layer in *C. hirsuta* stage 3 ovules (Fig. 1B, Fig. 6A). The cellular basis and 3D architecture of ii1’ layer formation is not well studied in non-Arabidopsis species, and therefore we explored if there are differences in the formation of this tissue between *C. hirsuta* and *A. thaliana*. In *C. hirsuta*, ii1’ initiation was observed from stage 3-I onward. Eight out of 10 stage 3-I ovules showed at least two ii1’ cells. Nine out of 10 stage 3-II ovules showed ii1’, and all ovules of stage 3-III up to stage 3-VI show the presence of ii1’. In stage 3-VI ovules, the ii1’ layer not only enveloped the proximal half of the inner integument, but also extended distally, covering more of the inner integument than in *A. thaliana*. In *C. hirsuta*, its distal border was also more irregular, with patches of cells separated from the proximal, more confluent ii1’ layer (Fig. 6A). We further noticed that the ii1’ layer in *C. hirsuta* occupied a significantly higher proportion of inner integument volume and cell number compared to that in *A. thaliana* (Fig. 6B,C). The growth of the ii1’ layer in terms of both tissue volume and cell number in *C. hirsuta* was double that in *A. thaliana*. Therefore, the ii1’ layer is disproportionately bigger in *C. hirsuta* compared to that in *A. thaliana*.

**Fig 6.**
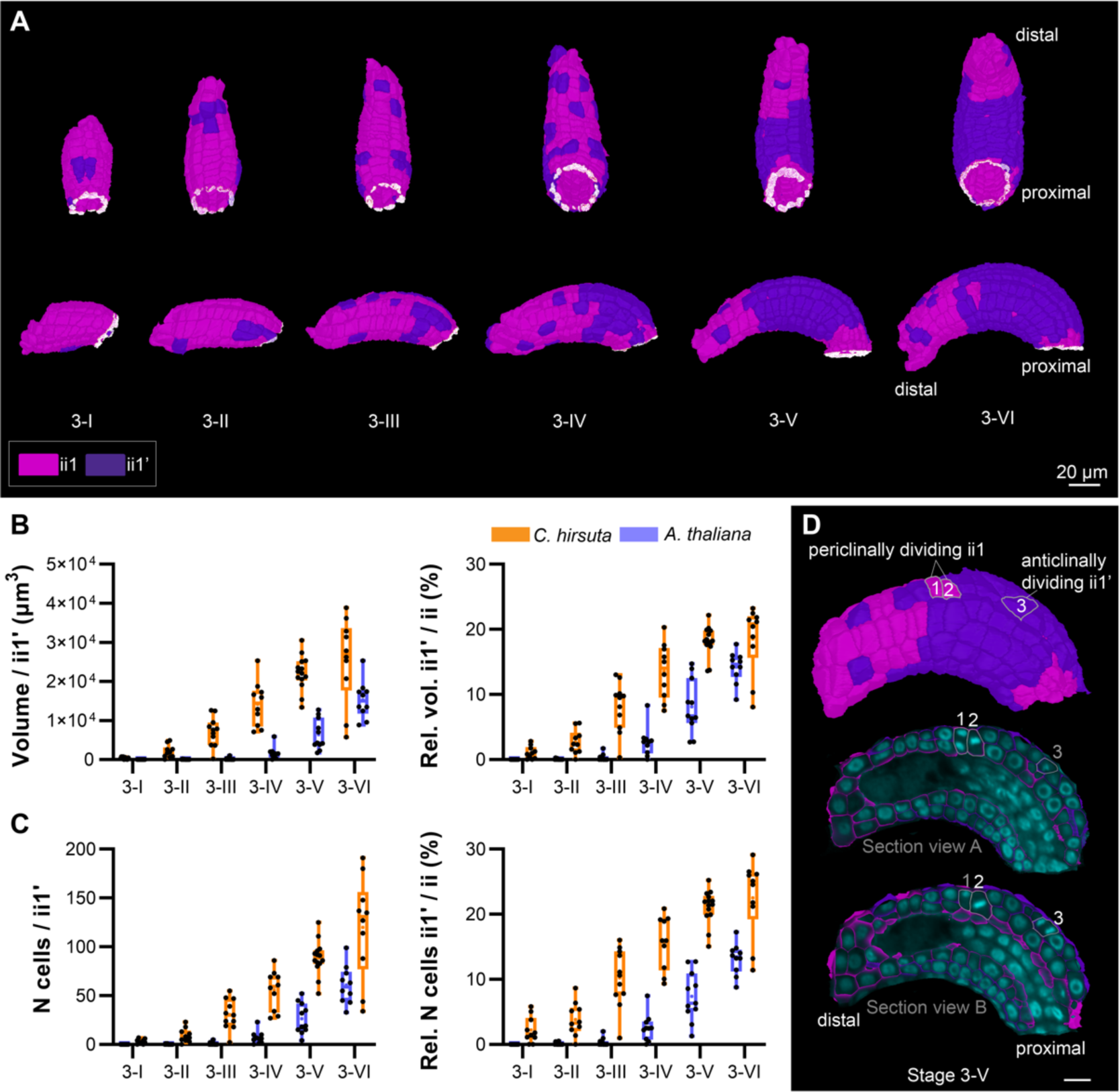
Development of parenchymatic inner integument layer in *C. hirsuta*. (A) The origin and propagation of the parenchymatic ii1 (ii1’) layer from stages 3-I up to 3-VI. Cells of the ii1’ layer are kept in blue. Upper panel: bottom view of 3D surface. Bottom panel: side view of 3D surface. Note that the ii1’ layer emerges as a patch of connected cells but later forms a ring-like structure of connected cells covering the proximal half of the inner integument, but still having patches of cells that are not a part of the ring. (B,C) Quantitative cellular analysis of the ii1’ layer. Data points indicate individual ovules. Mean ± SD is shown. Stages are indicated. (B) Left panel: plot depicting the total volume. Right panel: plot depicting the relative contribution of ii1’ volume to the volume of the entire inner integument. (C) Left panel: total cell number of ii1’. Right panel: the relative contribution of ii1’ total cell number to the cell number of the entire inner integument. (D) Side surface view (top) and section views (middle, bottom) of a stage 3-V ovule ii1 and ii1’ layers’ 3D cell mesh. In the section views, the overlaid nuclei z-stack showing the contribution of both periclinal divisions (cells 1 and 2 of ii1) and anticlinal divisions (cell 3 of ii1’) contribute to the propagation of *C. hirsuta* ii1’ layer. Scale bars: (A) 20 µm, (D) 10 µm.

To understand how this parenchymatous inner integument layer is initiated and propagated in *C. hirsuta*, we scored 3457 ii1’ cells in addition to all ii1 cells across the 62 ovules exhibiting an ii1’ layer, for the presence of mitotic divisions. Mitotic figures were visualized using the TO-PRO-3 nuclear stain (Fig. 6D). We observed 70 mitotic divisions in total. These were classified into periclinal ii1 divisions (40/70) and anticlinal ii1’ divisions (30/70). Therefore 57% of the mitotic divisions could be attributed to periclinal divisions of the ii1 cells. Thirty-four out of 40 (85 %) periclinal divisions were observed in stages 3-III to 3-VI. Twenty-five out of the 30 anticlinal divisions were observed in stages 3-V and 3-VI ovules; the remaining 5 belonged to stages 3-III and 3-IV. These results demonstrate that not only the initiation, but also the propagation of ii1’ in *C. hirsuta* involves periclinal ii1 cell divisions. Whereas, anticlinal divisions can only be attributed toward ii1’ propagation.

Taken together, these results suggest that the disproportionately larger growth of the *C. hirsuta* ii1’ cell layer is due not only to anticlinal cell divisions of ii1’ cells, but also to an approximately equal contribution of periclinal cell divisions in the ii1 layer. This is in contrast to ii1’ growth in *A. thaliana*, in which formation of the ii1’ layer involves a few scattered ii1 periclinal divisions, followed by ii1’ propagation mostly involving ii1’ anticlinal cell divisions (Vijayan et al., 2021).

## Discussion

Here, we provide a reference 3D digital atlas of *C. hirsuta* ovule development at full cellular resolution. First, the atlas provides a valuable resource for future genetic and molecular studies of ovule development in *C. hirsuta*. Second, it adds a high quality dataset to a growing collection of 3D digital organs that will be of considerable use not only for quantitative single-cell morphometric comparisons of 3D cellular architectures of ovules across plant species, but also for exploring the general properties of complex 3D cell assemblies. Third, the meshes can be used to generate templates for modeling ovule development. The original confocal z-stacks, segmented 3D images, meshes and analyses are available at the Biostudies database at EMBL-EBI (Sarkans et al., 2018) (accession number S-BIAD957).

A particular value of 3D digital organs with full cellular resolution is that they enable sophisticated quantitative analyses that allow the discovery of even subtle differences in cellular processes. Examples in *A. thaliana* include the control of division plane orientation in the embryo (Yoshida et al., 2014) and the regulation of 3D cell anisotropy in the hypocotyl (Montenegro-Johnson et al., 2015). In this study, it is exemplified by our comparative analysis of the growth patterns underlying the outgrowth of the ovule primordium and the development of the parenchymatous ii1’ layer of the inner integument between the two species *C. hirsuta* and *A. thaliana*. With respect to early ovule development, our results show that at the gross morphological level, primordium outgrowth in both species is similar and, except for size, results in primordia of comparable shape and layered structure. However, our data also suggest that the two species have distinct mechanisms for coordinating the relative contributions of the three radial layers to ovule primordium outgrowth. Similarity and diversity were also observed in the formation of the ii1’ layer. Both species develop this third layer of the inner integument. However, they vary mainly in the cellular processes that form the ii1’ layer, and also in the extent of this tissue. This indicates that the two species differ in the mechanisms that control the formation of the ii1’ layer. Taken together, these results strongly suggest that although morphogenetic processes may lead to organs of similar shapes, the underlying cellular processes can be different.

Topological analysis of *A. thaliana* 3D digital organs has been used to study organ design principles in plants (Bassel, 2019; Duran-Nebreda et al., 2023; Jackson et al., 2017; Jackson et al., 2019). For example, a recent network analysis of the diversity of cellular configurations in several morphologically simple 3D digital organs of *A. thaliana* suggested that the observed 3D cellular architectures at maturity are generated by active patterning mechanisms and do not result from random cell packing processes in these organs (Duran-Nebreda et al., 2023). Here, we applied topological analysis to the ovule, a plant organ of considerable complexity in terms of its 3D cellular architecture, tissue composition, and shape. Nerve-based analysis is a robust mathematical approach for studying how complex geometric objects are assembled from simple pieces (Edelsbrunner and Harer, 2010) that, to our knowledge, has not previously been applied to 3D digital organs. The goal was to identify differences in the 3D cellular organization of various tissues between the two species. We reasoned that mathematical analysis of cell connectivity maps could provide an approach for distinguishing cell populations that cannot be readily separated by visual inspection of morphology. In addition, it might provide independent confirmation of separate cell assemblies that previously could only be distinguished by morphological criteria, such as overall tissue size and shape or their 3D cellular architectures. As the nerve of a 3D digital ovule is a very complex structure, we compute feature vectors from the nerves that include only local information about the structure around each cell; this is a compromise between the desire to maintain as much information as possible, and the need for a representation of the data that is amenable to statistical analysis.

Interestingly, and despite the overall general similarity in shape between *C. hirsuta* and *A. thaliana* ovules, this approach led to the identification of clear topological differences in the 3D cellular architectures of the chalaza and integuments between the two species. This was especially surprising with respect to the outer integument, as visual inspection of its 3D cellular arrangements did not suggest obvious differences. Thus, nerve-based analysis represents a robust method to detect even apparently subtle distinctions in the topology of 3D digital organs with cellular resolution. The morphological consequences of such subtle differences in 3D cellular architectures remain to be determined. Our data indicate that *C. hirsuta* and *A. thaliana* ovules not only differ in a number of cellular parameters, size, and topology but also show subtle variations in their overall shape, such as the degree of curvature. Therefore, it will be interesting to investigate the relationship between the 3D cellular architecture of the ovule and its complex shape in future studies.

## Materials and Methods

### Plant work and lines

*C. hirsuta* var. Oxford (Ox), herbarium specimen voucher Hay 1 (OXF) (Hay and Tsiantis, 2006) and *Arabidopsis thaliana* (L.) Heynh. var. Columbia (Col-0) were used as wild-type strains. Plants were grown on soil as described earlier (Fulton et al., 2009; Hay et al., 2014).

### Clearing and staining of tissue samples

Treatment of *C. hirsuta* ovules was done as described in (Tofanelli et al., 2019) and (Vijayan et al., 2021) with some optimizations. Tissue was fixed in 4% paraformaldehyde in PBS for 1.5-2 h followed by one wash in PBS before transfer into the ClearSee solution (xylitol (10%, w/v), sodium deoxycholate (15%, w/v), urea (25%, w/v), in H_2_O) (Kurihara et al., 2015). Clearing was done at least overnight or for up to two to three days. Cell wall staining with SR2200 (Renaissance Chemicals, Selby, UK) was performed as described in (Musielak et al., 2015). Cleared tissue was washed in a PBS solution containing 0.1% SR2200 and then put into a PBS solution containing 0.1% SR2200 and a 1/1000 dilution of the nuclear stain TO-PRO-3 iodide (Thermo Fisher Scientific) for 30 minutes. Tissue was washed in PBS for one minute, transferred again to ClearSee for 20 minutes before mounting in Vectashield antifade agent (Vector Laboratories, Burlingame, CA, USA).

### Microscopy and image acquisition

Confocal laser scanning microscopy of ovules stained with SR2200 and TO-PRO-3 iodide was performed on an upright Leica TCS SP8 X WLL2 HyVolution 2 (Leica Microsystems) equipped with GaAsP (HyD) detectors and a 63x glycerol objective (HC PL anterior-posteriorO CS2 63x/1.30 GLYC, CORR CS2). Scan speed was at 400 Hz, the pinhole was set to 1 Airy units, line average between 2 and 4, and the digital zoom between 0.75 and 2. For z-stacks, 12 or 16 bit images were captured at a slice interval of 0.33 μm with voxel size of 0.126 μm x 0.126 μm x 0.33 μm. Laser power or gain was adjusted for z compensation to obtain an optimal z-stack. Image acquisition parameters were the following: SR2200 excitation, 405 nm diode laser (50 mW) with a laser power ranging from 0.1% to 1.5% intensity, detection at 416–476 nm with the gain of the HyD detector set to 20. TO-PRO-3 iodide excitation, white-light laser at 642 nm, with a laser power ranging from 2% to 3.5%, detection at 661 to 795 nm, with the gain of the HyD detector set to 400. Images were adjusted for color and contrast using ImageJ (Rueden et al., 2017) or MorphoGraphX software (Strauss et al., 2022).

### Datasets, 3D cell segmentation, and 3D cell meshes

The dataset encompassing the segmented wild-type 3D digital ovules of *A. thaliana* was described earlier (Vijayan et al., 2021). The z-stacks of *C. hirsuta* ovules were 3D cell segmented using the PlantSeg pipeline (Wolny et al., 2020). In all instances cell 3D meshes were generated with MorphoGraphX using segmented image stacks and the process “Mesh/Creation/Marching Cube 3D” with a cube size of 1. Manual cell type labeling was performed with MorphoGraphX.

### Exporting attributes from MorphoGraphX for further quantitative analysis

All quantitative cellular features were exported as attributes from MGX. The attributes included cell IDs (segmentation label of individual cells), cell type IDs (tissue annotation), and cell volume. The attributes from individual ovules were exported as csv files and merged to create long-format Excel-sheets listing all the scored attributes of all the cells from the analyzed ovules. Cell IDs with volume less than 30 μm^3^ have been excluded from cellular analyses since these correspond to artifacts and empty spaces that are segmented as cells. The files are included in the downloadable datasets.

### Topological analysis

#### The nerve construction

The nerve is closely related to the (unweighted) region adjacency graph, which has been used before to analyze 3D segmentations of plant tissues at cellular resolution (Wolny et al., 2020). This is the graph whose vertices are the cells, and whose edges are the pairs of cells that intersect (i.e., in the setting of 3D digital ovules, pairs of cells that share a voxel). To compute the nerve, one thinks of each cell as a vertex, and now computes every set of cells that intersect (i.e., a set *C*_1_, …, *C*_*k*_ of cells intersects if there is a voxel that belongs to *C*_i_ for all 1 ≤ *i* ≤ *k*). The edges of the region adjacency graph are exactly those sets of cells in the nerve that contain two cells. One pictures a set of two cells as an edge, a set of three cells as a triangle, and a set of four cells as a tetrahedron (see Fig. 4A); a set of cells with *k* elements is called a (*k* − 1)-simplex. Formally, the nerve is a *simplicial complex*, which is a generalization of a graph. When we analyze the cellular architecture of a particular tissue in the ovule, we compute the nerve of just those cells in that tissue.

#### Feature vectors

The feature vector of a nerve *N* is defined as follows. The *vertex star* of a vertex *v* of *N* is the set of *k*-simplices of *N* containing *v*, for all (≥ 0. The *face-vector* of *v* is the vector whose *k*-th component is the number of *k*-simplices in the vertex star of *v*. For example, in Fig. 4A, the vertex star of the red cell consists of one vertex, six edges, and six triangles, so its face-vector is (1,6,6). Note that the number of 1-simplices in the vertex star of *v* is exactly the number of neighbors of *v*, so this information is included in the face-vector. To define the feature vector *X* of *N*, we enumerate all possible face-vectors, then let the *i*-th component of *X* be the proportion of vertices of *N* whose face-vector is equal to the *i*-th face-vector in the enumeration. Note that, when we compute feature vectors for a set of nerves, it is necessary to enumerate all face-vectors vertices of all nerves, and use this enumeration when computing each feature vector.

#### Analysis of differences in topology

We analyzed the difference between species in the 3D cellular architecture of tissues during development in the following way. Fixing a tissue (e.g., the chalaza) and developmental stage, we computed the nerves of the cells of that tissue at that stage. This gives a set *N*_1_, …, *N*_*a*_ of nerves from *A. thaliana,* and a set *M*_1_, …, *M*_*b*_ of nerves from *C. hirsuta*. We computed the feature vectors of these nerves, giving a set *x*_1_, …, *X_a_* of *A. thaliana* feature vectors, and a set *Y*_1_, …, *Y_b_* of *C. hirsuta* feature vectors. We applied a multivariate two-sample test (see below) to the vectors *x*_1_, …, *X_a_* and *Y*_1_, …, *Y_b_* and computed the resulting p value. Looking at these p values across developmental stages gives insight into whether there is a difference between species in the 3D cellular architecture of the given tissue, and if so, at what stage in development this difference becomes visible in the nerve.

### Relative growth analysis

Relative growth between two consecutive stages was estimated by taking the ratio between the difference of the mean tissue volumes or the mean cell numbers of two consecutive stages and the corresponding mean parameter of the earlier stage according to the formula: (y(n+1) - y(n))/y(n).

### Software

The MorphoGraphX software was used for the generation of cell 3D meshes, cell type labeling, and the analysis of 3D cellular features (Barbier de Reuille et al., 2015; Strauss et al., 2022). It can be downloaded from its website (https://www.mpiz.mpg.de/MorphoGraphX). The PlantSeg pipeline (Wolny et al., 2020) was used for 3D cell boundary prediction and segmentation. The software can be obtained from its Github repository (https://github.com/hci-unihd/plant-seg). The C++ code, Python and R scripts, as well as all dependencies required for the topological analysis and its statistical evaluation were packaged using Docker (https://docker.com). The source code and the Dockerfiles can be obtained from the Github repository https://github.com/fabian-roll/NADO.

### Statistical analysis

Statistical analysis was performed using a combination of R (R Core Team, 2022) with RStudio (RStudio Team, 2020), the Anaconda distribution (Anaconda Software Distribution; https://anaconda.com) of the Python SciPy software stack (Oliphant, 2007), and PRISM10 software (GraphPad Software, San Diego, USA).

#### Box and whiskers plots

Boxplots show the median value of the distribution as a central line and mean value of the distribution as a plus sign within the box. The limits of the box represent the quartiles of the distribution. Whisker ends mark the minimum and maximum of all the data.

#### The multivariate two-sample test

To compare two sets of feature vectors, we use the multivariate two-sample test of Baringhaus and Franz (Baringhaus and Franz, 2004). Given independent random vectors in d-dimensional Euclidean space X1, …, Xm and Y1, …, Yn that are identically distributed with distribution functions F and G, one tests the null hypothesis F = G against the general alternative F ≠ G using a test statistic called the Cramér statistic that is defined using Euclidean distances. Distances between Xs and Ys contribute to the statistic with positive weight, while distances between Xs and distances between Ys contribute with negative weight. Rejection of the null hypothesis is for large positive values of the test statistic. Baringhaus and Franz show that p values can be estimated using bootstrapping (Van der Vaart and Wellner, 2012). The estimated p value can be zero, if the computed value of the test statistic is larger than all the values computed from bootstrap samples. The test is implemented by the R package “cramer” (Franz, 2019).

## Acknowledgements

We thank Angela Hay and Miltos Tsiantis for providing seeds of wild-type *C. hirsuta*. We further thank Gabriella Mosca and members of Schneitz lab for comments on the manuscript. We acknowledge support by the Center for Advanced Light Microscopy (CALM) of the TUM School of Life Sciences.

## Data availability

The *C. hirsuta* 3D digital ovule dataset has been deposited with the BioStudies data repository at EMBL-EBI (Sarkans et al., 2018) (https://www.ebi.ac.uk/biostudies) (accession number S-BIAD957). It includes raw cell boundaries, cell boundaries, PlantSeg predictions, nuclei images, segmented cells, and the annotated 3D cell meshes along with the associated attribute files in csv format. The 3D mesh files can be opened in MorphoGraphX. It also includes the *C. hirsuta* long format cell attributes files and the data files used for topological analysis. The corresponding Arabidopsis ovule dataset on Biostudies has the accession numbers S-BSST475 (raw and topological analysis data) and S-BSST513 (long format cell attribute file).

## Authors’ contributions

TAM and KS designed the study. TAM generated the Cardamine atlas and performed the morphology experiments. TAM and KS interpreted the results. AR, NS, FR, and UB conceived, performed, analyzed, and interpreted the topological analysis. KS and UB secured funding. TAM, AR, FR, NS, and KS wrote the paper. All authors read and approved the final manuscript.

---

**Figure S1:**
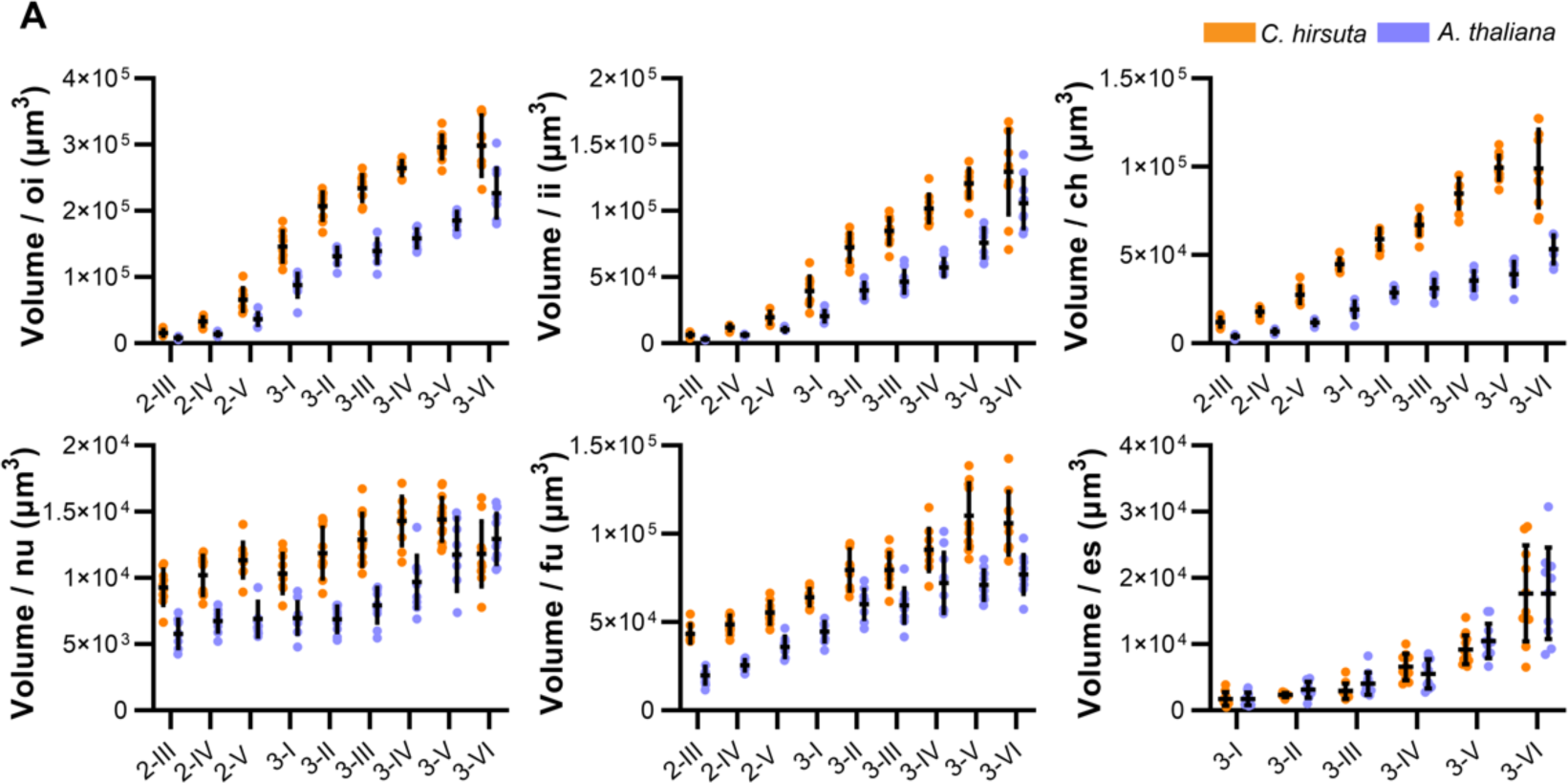
Comparison of tissue-specific volumes of *C. hirsuta* and *A. thaliana* ovules. Plots comparing total tissue volumes of the outer integument, inner integument, chalaza, nucellus, funiculus, and embryo sac of wild type *C. hirsuta* and *A. thaliana* ovules at stages 2-III to 3-VI. Data points indicate individual ovules. Mean ± SD are represented as bars and whiskers. Abbreviations: ch, chalaza; es, embryo sac; fu, funiculus; ii, inner integument; nu, nucellus; oi, outer integument.

**Figure S2.**
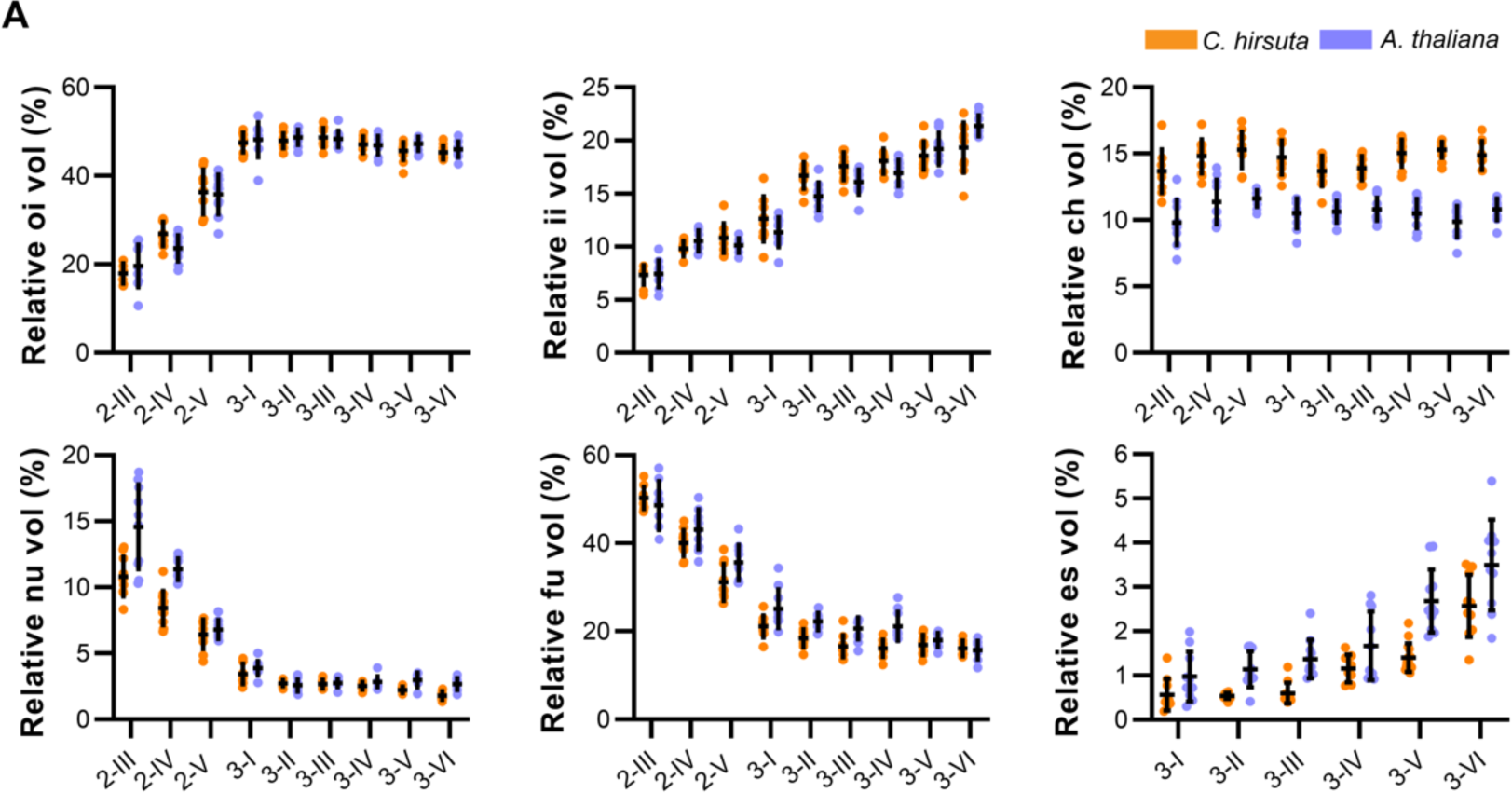
Comparative differences in relative contributions of tissue-specific volumes to the entire ovule. Plots comparing the relative contribution of the volume of each tissue to that of the entire ovule for both *C. hirsuta* and *A. thaliana* at different stages. Data points indicate individual ovules. Mean ± SD are represented as bars and whiskers. Abbreviations: ch, chalaza; es, embryo sac; fu, funiculus; ii, inner integument; nu, nucellus; oi, outer integument.

**Figure S3.**
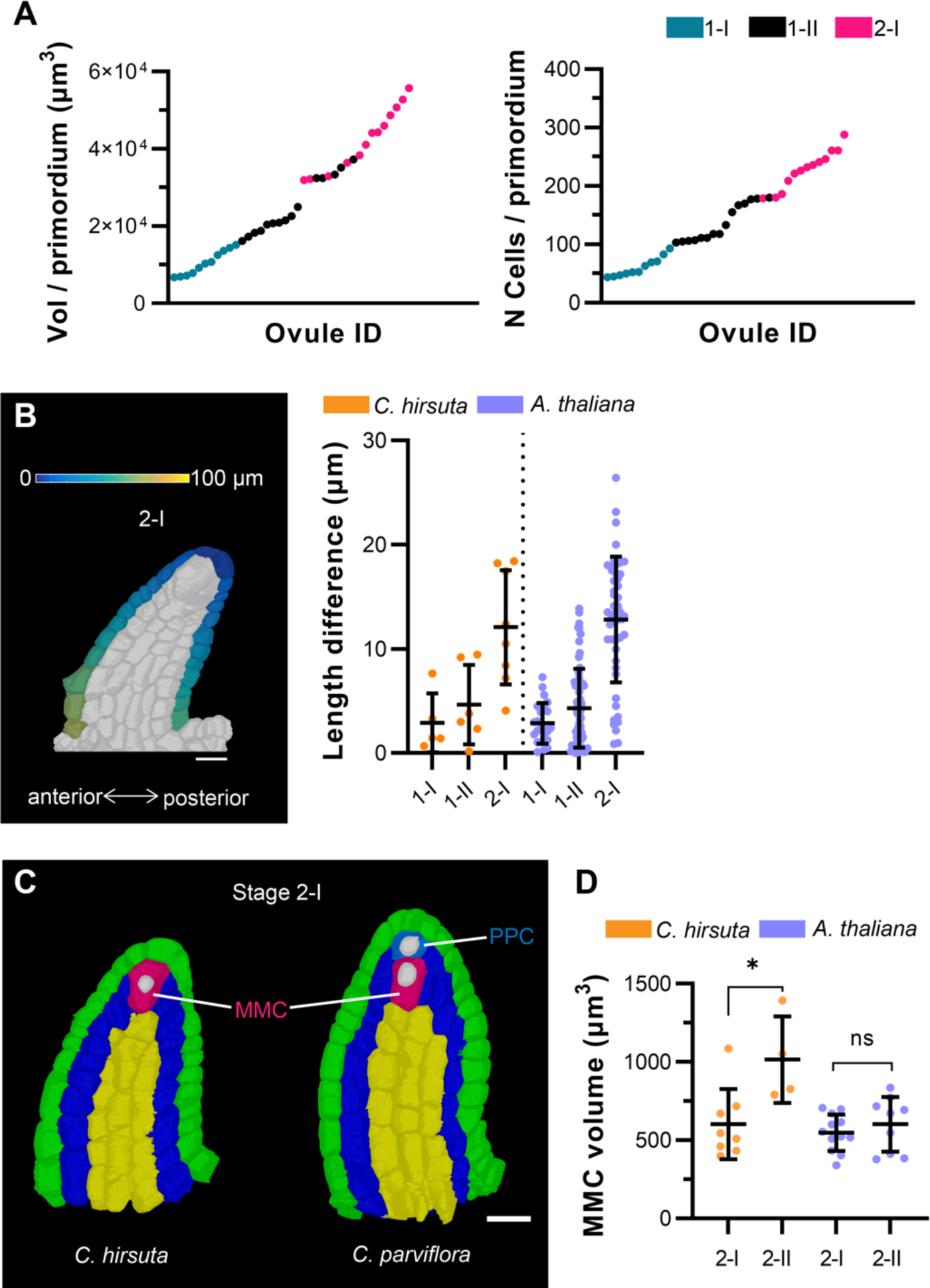
Developmental features underlying the growth of *C. hirsuta* ovule primordia. (A) *C. hirsuta* primordia grow in a continuous manner. (A) Left panel: plot indicating the total volume of primordia ordered according to increasing volume. Right panel: plot depicting the total number of cells in the ovules ordered according to the increasing number of cells. Data points indicate individual ovules and are colored by stage. (B) C. hirsuta primordia show slanting. Left panel: 2D section view of a stage 2-I 3D cell mesh; the heatmap on the surface cells of posterior and anterior halves depicts the quantified distance value between individual measured cells to the distal tip of primordia. Right panel: plot showing a comparison of the extent of slanting of *C. hirsuta* and *A. thaliana* primordia, quantified by the difference in maximal length on the anterior and posterior sides of ovule at stages 1-I, 1-II, and 2-I. Data points indicate individual ovules. Mean ± SD are represented as bars and whiskers. (C) MMC development in Cardamine. *C. hirsuta* is tenuinucellate; the MMC is directly located below the epidermis. *C. parviflora* is crassinucellate; the MMC is located below the primary parietal cell that lies between the epidermis and MMC. Scale bar 10 µm. (D) Plot showing a comparison of MMC volume of stage 2-I and 2-II primordia of *C. hirsuta* and *A. thaliana*. Mean ± SD are represented as bars and whiskers. Abbreviations: MMC, megaspore mother cell; PPC, primary parietal cell.

**Table S1.**
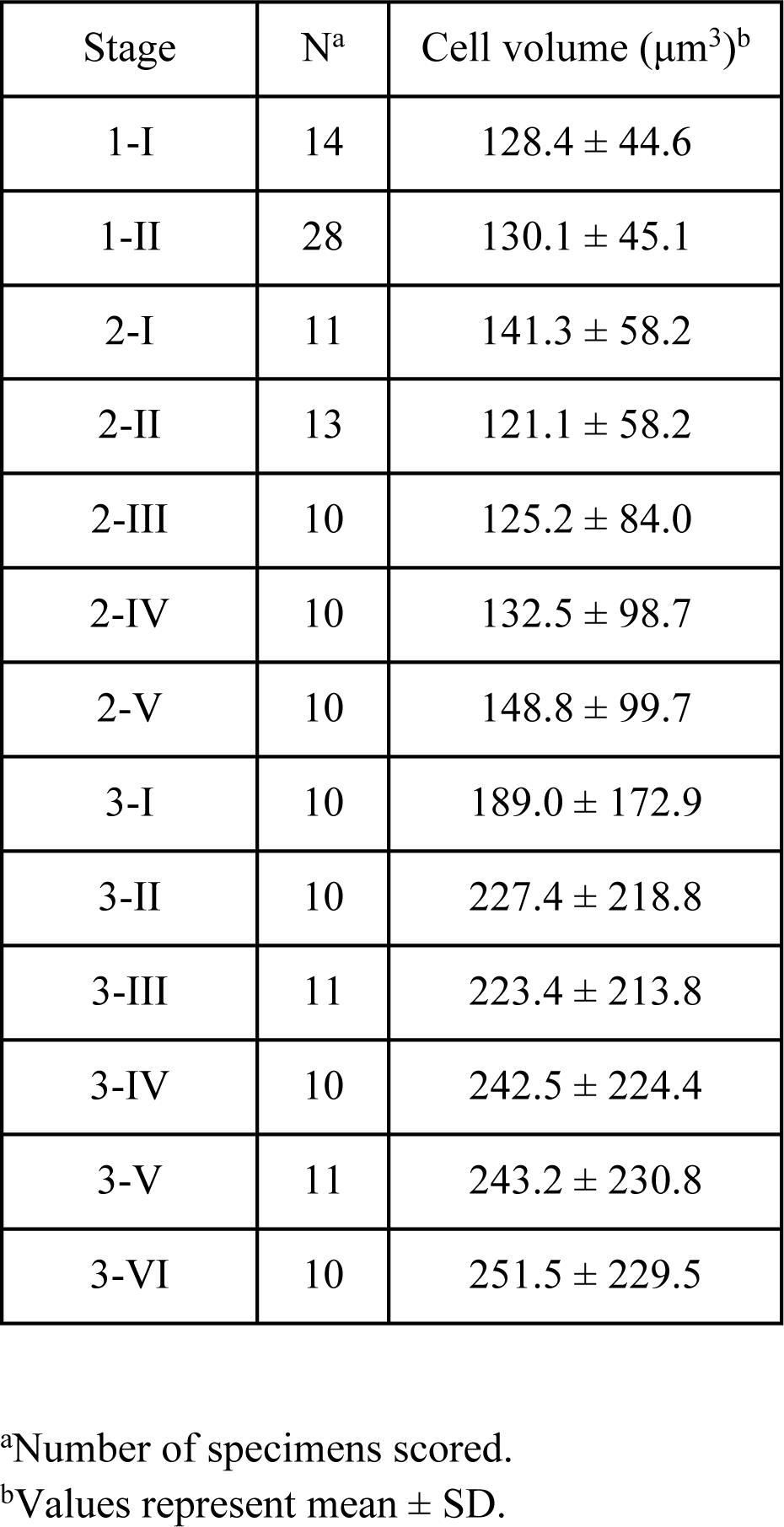
Cell volumes of *A. thaliana* ovules at different developmental stages.

**Table S2.**
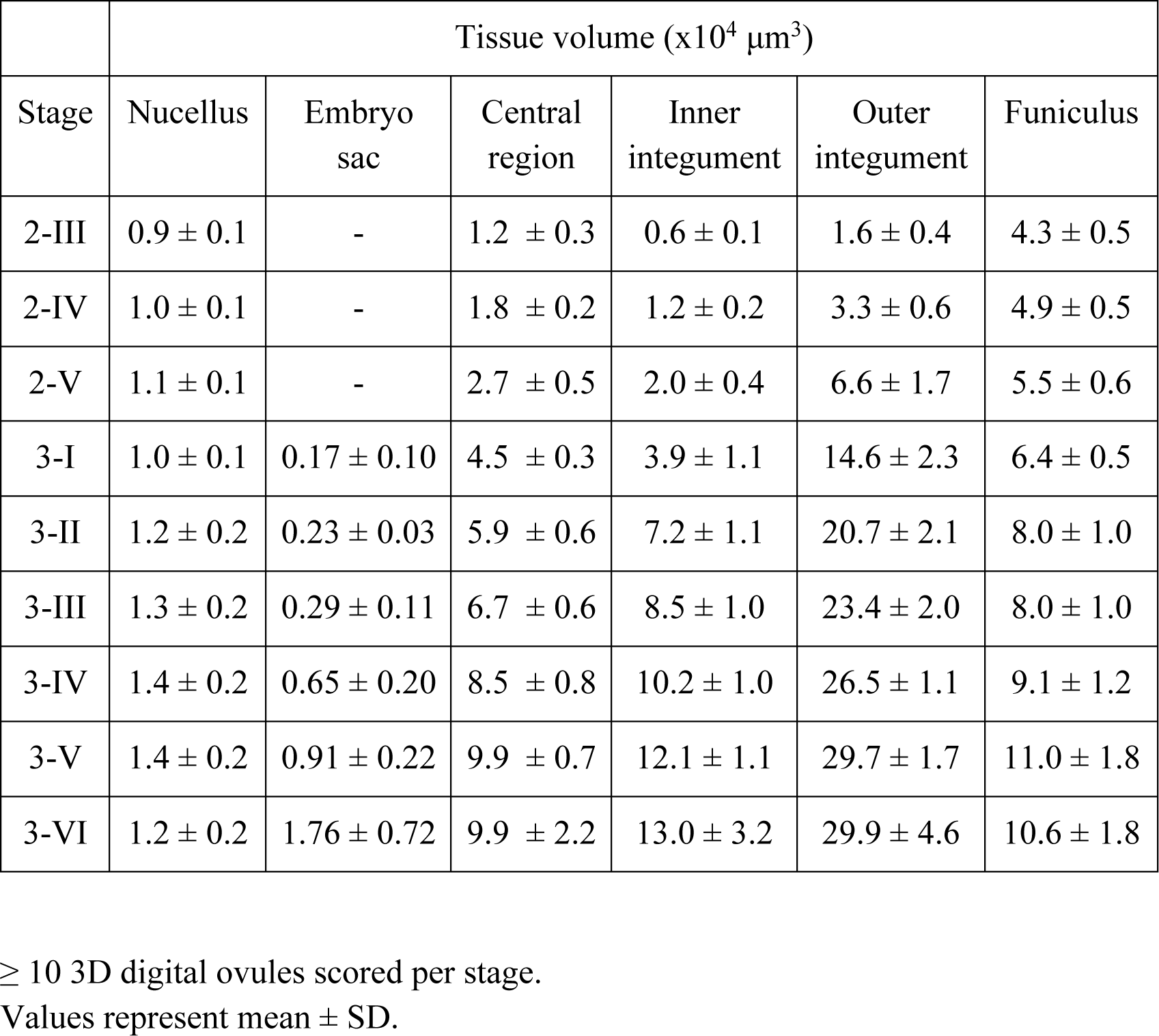
Total volumes of major ovule tissues in *C. hirsuta*.

|  | Tissue volume (x10<sup>4</sup> μm<sup>3</sup>) |  |  |  |  |  |
| --- | --- | --- | --- | --- | --- | --- |
| Stage | Nucellus | Embryo sac | Central region | Inner integument | Outer integument | Funiculus |
| 2-III | 0.9 ± 0.1 | - | 1.2 ± 0.3 | 0.6 ± 0.1 | 1.6 ± 0.4 | 4.3 ± 0.5 |
| 2-IV | 1.0 ± 0.1 | - | 1.8 ± 0.2 | 1.2 ± 0.2 | 3.3 ± 0.6 | 4.9 ± 0.5 |
| 2-V | 1.1 ± 0.1 | - | 2.7 ± 0.5 | 2.0 ± 0.4 | 6.6 ± 1.7 | 5.5 ± 0.6 |
| 3-I | 1.0 ± 0.1 | 0.17 ± 0.10 | 4.5 ± 0.3 | 3.9 ± 1.1 | 14.6 ± 2.3 | 6.4 ± 0.5 |
| 3-II | 1.2 ± 0.2 | 0.23 ± 0.03 | 5.9 ± 0.6 | 7.2 ± 1.1 | 20.7 ± 2.1 | 8.0 ± 1.0 |
| 3-III | 1.3 ± 0.2 | 0.29 ± 0.11 | 6.7 ± 0.6 | 8.5 ± 1.0 | 23.4 ± 2.0 | 8.0 ± 1.0 |
| 3-IV | 1.4 ± 0.2 | 0.65 ± 0.20 | 8.5 ± 0.8 | 10.2 ± 1.0 | 26.5 ± 1.1 | 9.1 ± 1.2 |
| 3-V | 1.4 ± 0.2 | 0.91 ± 0.22 | 9.9 ± 0.7 | 12.1 ± 1.1 | 29.7 ± 1.7 | 11.0 ± 1.8 |
| 3-VI | 1.2 ± 0.2 | 1.76 ± 0.72 | 9.9 ± 2.2 | 13.0 ± 3.2 | 29.9 ± 4.6 | 10.6 ± 1.8 |

≥ 10 3D digital ovules scored per stage.
Values represent mean ± SD.

**Table S3.**
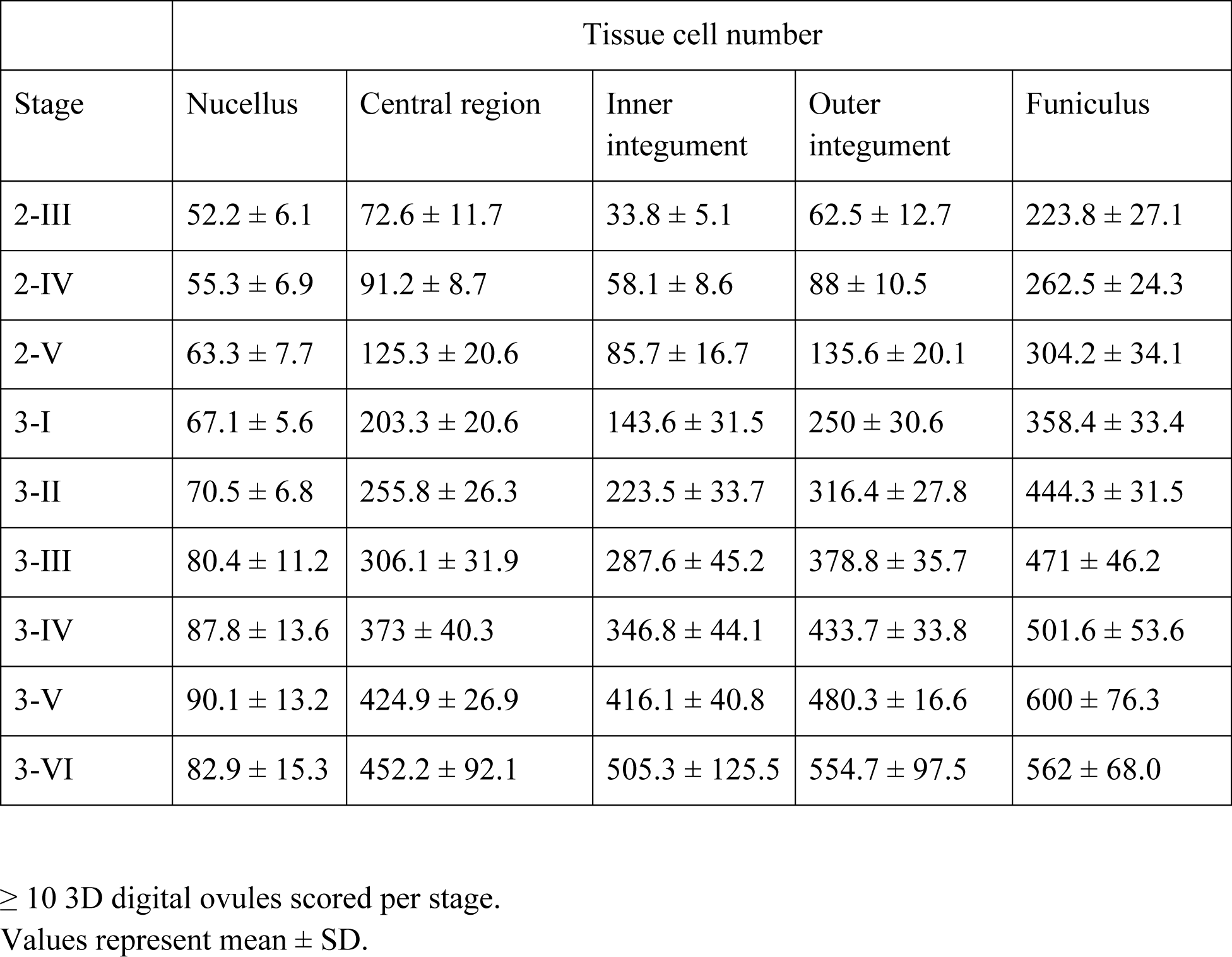
Total cell numbers of major ovule tissues in *C. hirsuta*.

|  | Tissue cell number |  |  |  |  |
| --- | --- | --- | --- | --- | --- |
| Stage | Nucellus | Central region | Inner integument | Outer integument | Funiculus |
| 2-III | 52.2 ± 6.1 | 72.6 ± 11.7 | 33.8 ± 5.1 | 62.5 ± 12.7 | 223.8 ± 27.1 |
| 2-IV | 55.3 ± 6.9 | 91.2 ± 8.7 | 58.1 ± 8.6 | 88 ± 10.5 | 262.5 ± 24.3 |
| 2-V | 63.3 ± 7.7 | 125.3 ± 20.6 | 85.7 ± 16.7 | 135.6 ± 20.1 | 304.2 ± 34.1 |
| 3-I | 67.1 ± 5.6 | 203.3 ± 20.6 | 143.6 ± 31.5 | 250 ± 30.6 | 358.4 ± 33.4 |
| 3-II | 70.5 ± 6.8 | 255.8 ± 26.3 | 223.5 ± 33.7 | 316.4 ± 27.8 | 444.3 ± 31.5 |
| 3-III | 80.4 ± 11.2 | 306.1 ± 31.9 | 287.6 ± 45.2 | 378.8 ± 35.7 | 471 ± 46.2 |
| 3-IV | 87.8 ± 13.6 | 373 ± 40.3 | 346.8 ± 44.1 | 433.7 ± 33.8 | 501.6 ± 53.6 |
| 3-V | 90.1 ± 13.2 | 424.9 ± 26.9 | 416.1 ± 40.8 | 480.3 ± 16.6 | 600 ± 76.3 |
| 3-VI | 82.9 ± 15.3 | 452.2 ± 92.1 | 505.3 ± 125.5 | 554.7 ± 97.5 | 562 ± 68.0 |

≥ 10 3D digital ovules scored per stage.
Values represent mean ± SD.

**Table S4.**
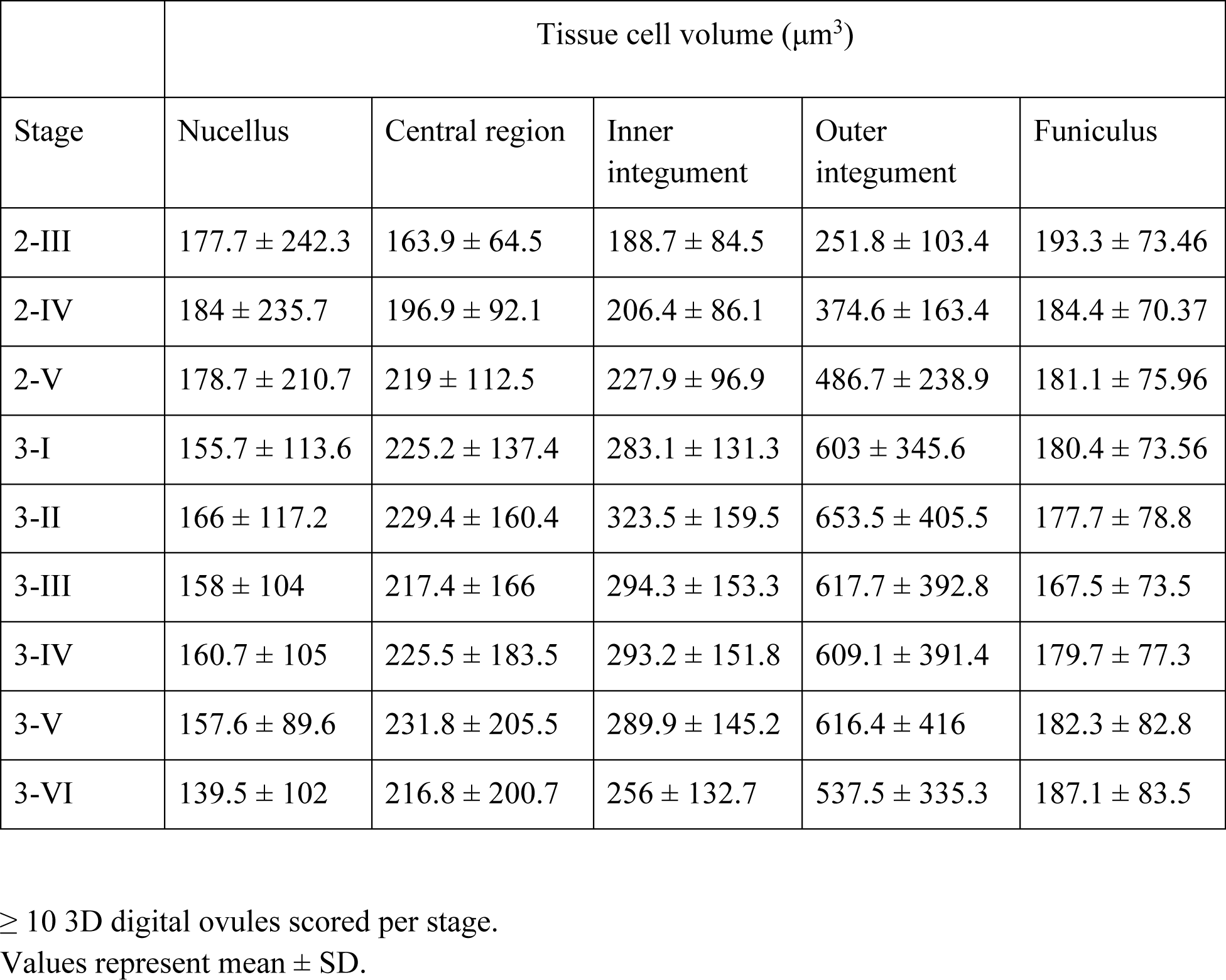
Cell volumes of major ovule tissues in *C. hirsuta*.

**Table S5.** P values from two-sample test applied to nerve-based feature vectors of *A. thaliana* and *C. hirsuta*.

| Stage | Ovule | Outer integument | Inner integument | Chalaza | Nucellus | Funiculus |
| --- | --- | --- | --- | --- | --- | --- |
| 2-III | $1.1 \times 10^{-3}$ | $4.6 \times 10^{-2}$ | $1.3 \times 10^{-2}$ | $9.9 \times 10^{-2}$ | $1.6 \times 10^{-1}$ | $1.5 \times 10^{-1}$ |
| 2-IV | 0 | $1.3 \times 10^{-2}$ | $1.7 \times 10^{-1}$ | $2.0 \times 10^{-1}$ | $1.3 \times 10^{-1}$ | $9.0 \times 10^{-2}$ |
| 2-V | $1.8 \times 10^{-2}$ | $8.4 \times 10^{-2}$ | $3.7 \times 10^{-1}$ | $2.0 \times 10^{-1}$ | $4.6 \times 10^{-1}$ | $1.4 \times 10^{-1}$ |
| 3-I | $5.0 \times 10^{-4}$ | $7.0 \times 10^{-4}$ | $3.2 \times 10^{-1}$ | $2.0 \times 10^{-2}$ | $3.3 \times 10^{-1}$ | $1.2 \times 10^{-2}$ |
| 3-II | 0 | $6.5 \times 10^{-3}$ | $1.6 \times 10^{-1}$ | $9.5 \times 10^{-3}$ | $7.1 \times 10^{-2}$ | $3.0 \times 10^{-2}$ |
| 3-III | $1.0 \times 10^{-4}$ | $7.5 \times 10^{-3}$ | $1.1 \times 10^{-3}$ | $2.0 \times 10^{-4}$ | $8.0 \times 10^{-2}$ | $7.5 \times 10^{-2}$ |
| 3-IV | $8.3 \times 10^{-2}$ | $1.0 \times 10^{-3}$ | $4.0 \times 10^{-4}$ | $9.6 \times 10^{-3}$ | $2.5 \times 10^{-1}$ | $1.9 \times 10^{-1}$ |
| 3-V | 0 | $1.0 \times 10^{-4}$ | $1.0 \times 10^{-4}$ | $1.0 \times 10^{-4}$ | $1.3 \times 10^{-1}$ | $1.3 \times 10^{-2}$ |
| 3-VI | $9.0 \times 10^{-4}$ | $6.0 \times 10^{-4}$ | $5.2 \times 10^{-3}$ | $9.0 \times 10^{-4}$ | $2.8 \times 10^{-1}$ | $5.6 \times 10^{-2}$ |

